# A core speech circuit between primary motor, somatosensory, and auditory cortex: evidence from connectivity and genetic descriptions

**DOI:** 10.1101/139550

**Authors:** Jeremy I. Skipper, Uri Hasson

**Author notes:** Address correspondence to: Jeremy I Skipper, University College London, Experimental Psychology, 26 Bedford Way, London WC1H OAP, United Kingdom.

## Abstract

What adaptations allow humans to produce and perceive speech so effortlessly? We show that speech is supported by a largely undocumented core of structural and functional connectivity between the central sulcus (CS or primary motor and somatosensory cortex) and the transverse temporal gyrus (TTG or primary auditory cortex). Anatomically, we show that CS and TTG cortical thickness covary across individuals and that they are connected by white matter tracts. Neuroimaging network analyses confirm the functional relevance and specificity of these structural relationships. Specifically, the CS and TTG are functionally connected at rest, during natural audiovisual speech perception, and are coactive over a large variety of linguistic stimuli and tasks. Importantly, across structural and functional analyses, connectivity of regions immediately adjacent to the TTG are with premotor and prefrontal regions rather than the CS. Finally, we show that this structural/functional CS-TTG relationship is mediated by a constellation of genes associated with vocal learning and disorders of efference copy. We propose that this core circuit constitutes an interface for rapidly exchanging articulatory and acoustic information and discuss implications for current models of speech.

---

> “… for as everyone knows, parrots can talk.”
>
> — Darwin (1871, p. 54)

## 1 Introduction

Some animals like parrots and humans are highly adept at vocal learning and perceiving complex auditory sequences. Others vary on a continuum from non-learning animals like chickens, to more limited or moderate vocal learners, like mice and monkeys, to complex vocal and sequence learning species like songbirds (Petkov & Jarvis, 2012). What neurobiological changes resulted in these differences in skill across animals? To address this question, we review some comparative brain anatomy, which suggests that the number and distribution of neuronal connections has significantly changed in the human brain. We propose that these new connections might allow for greater sensorimotor flexibility by providing a pathway for “efference copy” signals. These are copies of motor commands from late in the motor pathway to early in the auditory sensory pathway that support vocal learning and speech production and perception by allowing for comparison of actual with intended/target acoustic patterns. Consistent with this proposal, emerging and mostly indirect evidence suggests that primary motor, primary somatosensory, and primary auditory regions are often coactive and, thus, might provide a substrate for efference copy and the human adeptness with speech. After this review, we introduce a set of six studies that explicitly address this suggestion. Namely, we test the hypotheses that this set of regions form a “core” of structural and functional connectivity and that activity in this network can be speech specific, both with regard to functional properties and associated genetic phenotype.

### 1.1 Comparative anatomy

To understand what makes humans so dexterous with speech, a seemingly good place to look would be other primates who have speech-ready vocal tracts yet do not talk (Tecumseh Fitch, de Boer, Mathur, & Ghazanfar, 2016). Vocal readiness and limited vocal learning in non-human primates, considered together with human brain expansion, all suggest that differences are neurobiological in origin. One logical difference might be in the extent of brain connectivity between areas associated with speech. The arcuate fasciculus white-matter tract, connecting frontal, parietal, and temporal regions, has long been associated with uniting “Wernicke’s” and “Broca’s” language areas (e.g., Geschwind, 1970). Indeed, this tract has significantly expanded in the number of temporal regions it contacts in humans compared to other primates (Rilling et al., 2008; Rilling, Glasser, Jbabdi, Andersson, & Preuss, 2011).

Another set of significantly-altered connections are those between laryngeal motor cortex and the brainstem and other cortical regions. Whereas nonhuman primate laryngeal motor cortex is in premotor cortex and is thought to have only indirect projections to brainstem neurons, human laryngeal motor cortex has direct connections (Brown, Ngan, & Liotti, 2008; Simonyan, 2014; Simonyan & Horwitz, 2011). In addition to these changes in subcortical connections, the distribution of laryngeal motor cortical connections has also changed. The majority of nonhuman primate laryngeal motor cortex structural connections are with other frontal regions and the putamen. In contrast, human laryngeal motor cortex has proportionally less connections to these regions and about a four-fold increase in primary somatosensory and nine-fold increase in inferior parietal lobule connectivity (Kumar, Croxson, & Simonyan, 2016). In addition, human laryngeal motor cortex has a greater spread of connectivity with superior temporal auditory regions (Kumar et al., 2016).

These differences suggest that more complex vocal learning phenotypes of the sort seen in humans rely on connectivity patterns that provide more flexible control over vocalizations. That is, direct control of brainstem laryngeal motoneurons by laryngeal motor cortex would seem to permit more voluntary control over vocalizations. Increased frontal motor, somatosensory, and auditory connectivity might provide a substrate by which animals could *learn* to flexibly vocalize. In contrast, limited vocal learners like monkeys, without this pattern of connectivity, rely more on subcortical and medial cortical connections and produce less flexible and more involuntary or affective sounds (Petkov & Jarvis, 2012; Simonyan & Horwitz, 2011).

### 1.2 Efference copy

How does greater motor, somatosensory, and auditory connectivity translate into vocal flexibility? One possibility is through efference copies. These are copies of the late efferent vocal motor commands issued to early sensory cortices (Crapse & Sommer, 2008). Efference copies correspond to the *predicted* sensory targets of a vocalization that can be compared to actual sensory feedback. A difference is used to adjust vocalizations to better match a target sound during learning or make real-time corrections to perturbations of the articulators. Indeed, neuroimaging and neurophysiological studies suggest that efference copy plays an important role in human vocal learning and speech production (Brown, Ingham, Ingham, Laird, & Fox, 2005; Chang, Niziolek, Knight, Nagarajan, & Houde, 2013; C.-M. A. Chen et al., 2011; Guenther, Ghosh, & Tourville, 2006; Kingyon et al., 2015; Niziolek, Nagarajan, & Houde, 2013; Rummell, Klee, & Sigurdsson, 2016; J. Wang et al., 2014).

Reuse (Anderson, 2010) or recycling (Dehaene & Cohen, 2007) of this efference copy mechanism might also help explain how we perceive speech. That is, it is still unknown how we achieve perceptual constancy given how variable the acoustic patterns reaching our cochleas are. This indeterminacy between physical signals and how they are perceived might be resolved by knowledge based predictions from vocal motor systems (Jeremy I. Skipper, Devlin, & Lametti, 2017; J. I. Skipper, Nusbaum, & Small, 2006). For example, hearing “we went down to the pond to listen to the croaking…” facilitates access to words like “frogs”. These can be sequenced by the vocal motor system resulting in activation of motor programs for saying /f/, /r/, etc. An /f/ efference copy from motor to auditory cortex can constrain interpretation of the subsequent patterns of activity arriving in auditory cortex by serving as an “hypothesis” about what pattern is expected (Jeremy I. Skipper & Zevin, 2017). Indeed, neuroimaging studies suggest that predictive or efference copy like mechanisms support speech perception (Dikker & Pylkkänen, 2012; Dikker, Silbert, Hasson, & Zevin, 2014; Ettinger, Linzen, & Marantz, 2014; Frank & Willems, n.d.; Gambi & Pickering, 2013; Leonard, Bouchard, Tang, & Chang, 2015; Pascale Tremblay, Deschamps, Baroni, & Hasson, 2016; Willems, Frank, Nijhof, Hagoort, & van den Bosch, 2015) and that these predictions involve speech production systems (Jeremy I. Skipper, 2014; Jeremy I. Skipper et al., 2017; Jeremy I. Skipper, van Wassenhove, Nusbaum, & Small, 2007; Jeremy I. Skipper & Zevin, 2017).

### 1.3 Neurobiological models

Thus, human adeptness with speech production and perception may be supported by increased cortical and subcortical connectivity, permitting increased flexibility with vocalizations via a predictive efference copy mechanism. Consistent with this, classical (Geschwind, 1970) and most contemporary models of the neurobiology of speech (Friederici, 2011; Hickok, Houde, & Rong, 2011; Hickok & Poeppel, 2007; Rauschecker & Scott, 2009) propose functional links between frontal and temporal regions. These models, however, largely focus on interfaces between auditory *association* cortices and what are considered “high-level” speech-related systems. As such, the relationship between the anterior auditory association cortex and mostly prefrontal regions is well described in humans and other primates (Romanski & Averbeck, 2009). Similarly, the relationship between posterior auditory association cortex (like the planum temporale) and higher level motor regions like the pars opercularis, and dorsal and ventral premotor cortex is also well described (e.g., Hickok, 2012b).

However, in contrast to the higher level or association regions described in current models of speech, efference copy, as alluded to earlier, is often suggested to reflect a “low-level” process across phylum (Crapse & Sommer, 2008; Griisser, 1995). The term low-level, as used here, refers to interactions between motor regions closer to innervating muscles and auditory regions closer to the auditory periphery. Indeed, motor-auditory efference copy is well documented in crickets (Poulet & Hedwig, 2003, 2006, 2007) and vocal fish (Chagnaud & Bass, 2013), animals that presumably have more rudimentary motor or auditory systems. Mice, despite not being highly skilled vocal learners, show evidence for efference copy through connectivity between motor and early auditory cortices (Nelson et al., 2013; Schneider & Mooney, 2015; Schneider, Nelson, & Mooney, 2014). Likewise, auditory cortex in marmosets, a vocal primate, has neurophysiological properties consistent with efference copy during vocalization though the pathway through which this modulation occurs is unknown (S. J. Eliades & Wang, 2012; Steven J. Eliades & Wang, 2002, 2008, 2013; Ghazanfar & Eliades, 2014). Finally, in vocal learning songbirds, efference copy between motor and auditory regions has been implicated in both online and offline (i.e., sleep related) vocal learning, vocal error correction, song maintenance, and listening (Brainard & Doupe, 2000; Dave & Margoliash, 2000; Margoliash, 2002; Schneider & Mooney, 2015; Sober & Brainard, 2009; Troyer & Doupe, 2000; Tumer & Brainard, 2007).

### 1.4 A speech core?

Interactions between *lower-level motor and auditory regions* have not been extensively considered or empirically demonstrated (as we review below). For this reason, they are not integrated into mainstream models of human speech perception (like popular “dual-stream” models)(Hickok & Poeppel, 2007; Rauschecker & Scott, 2009). We suggest it is time to reconsider this position given the animal literature discussed above and a body of recent evidence demonstrating that these lower-level audiomotor regions do play an important role in speech production and perception. In particular, Markiewicz and Bohland (2016) demonstrated coding of syllable identity in ventral CS during speech production. Evans and Davis (2015) documented acoustically-invariant responses to syllable identity in ventral CS during speech perception. Further suggestive of an efference copy mechanism, Liebenthal et al. (2013) used simultaneously acquired EEG and fMRI and found that the ventral CS, inferior parietal, and superior temporal regions were strongly correlated as early as 100 ms after speech stimulus onset suggesting “*efferent* feedback from somatomotor to auditory cortex” (pg. 15,419; our emphasis).

Consistent with this account, efference copy and predictive models propose a reduction in activity when auditory stimuli are more predictable (e.g., due to reduced magnitudes of prediction error). Indeed, both the CS and TTG show reduced activity when auditory and linguistic predictability increases during perception (Jeremy I. Skipper, 2014; Pascale Tremblay et al., 2016). Finally, these neuroimaging studies are supported by an independent body of work using transcranial magnetic stimulation that suggests a causal involvement of primary motor cortex in speech perception (Möttönen & Watkins, 2009; Schomers, Kirilina, Weigand, Bajbouj, & Pulvermüller, 2014; Smalle, Rogers, & Möttönen, 2014).

In addition to these studies, several recent nonhuman and human primate studies indirectly suggest a strong relationship between primary motor, somatosensory, and auditory cortex despite not being discussed as such in that work. In monkeys, stimulating the mid/caudal lateral belt, adjacent to primary auditory cortex, produces strong activation of the CS (Petkov et al., 2015). Furthermore, it seems that humans have significantly thicker and more gyrification in the CS and TTG than chimpanzees (Hopkins, Li, Crow, & Roberts, 2016). Human CS and TTG are the most highly myelinated regions in the human brain (Carey, Krishnan, Callaghan, Sereno, & Dick, 2017; Lutti, Dick, Sereno, & Weiskopf, 2014) and may be connected by white matter tracts (Fan et al., 2016). These structural differences are paralleled by task-free or “resting-state” (Jorge Sepulcre, 2015; J. Sepulcre, Sabuncu, Yeo, Liu, & Johnson, 2012; Yeo et al., 2011) and task-based functional connectivity (Krienen, Yeo, & Buckner, 2014; Laird et al., 2011; Smith et al., 2009). Finally, the CS and TTG evince functional network-by-gene interactions (Krienen, Yeo, Ge, Buckner, & Sherwood, 2016).

### 1.5 Hypotheses

To conclude, new findings suggesting the involvement of the CS in representing auditory, phonetic, and syllabic level information have not been integrated within current neurobiological models of speech and language, but this and other less direct evidence suggest a “lower-level” circuit between primary motor, somatosensory, and auditory regions. This interface might constitute a backbone or core circuit for vocal learning, speech production, and speech perception through a rapid efference copy mechanism between these regions. Our primary hypothesis in the current work is that, if this circuit exists in humans, then it should be accompanied by anatomically specific structural and functional connectivity between the central sulcus (CS) or Brodmann’s cytoarchitectonic area (BA) 4 and 3, encompassing primary motor and primary somatosensory cortices and the transverse temporal gyrus (TTG) or roughly BA 41/42, encompassing primary auditory cortex. This connectivity should be qualitatively different form the connections of adjacent secondary auditory regions, including the immediately adjacent planum polare, planum temporale, and superior temporal gyrus (STG). Furthermore, these connections should be in evidence across multiple signatures of structural and functional connectivity, with the latter demonstrating functional properties consistent with a role in speech, vocal learning, and efference copy.

To evaluate these hypotheses, we conducted six neuroimaging analyses, including two structural and three functional connectivity and one network-by-gene description analysis. First, we analyzed cortical thickness data using structural MRI to test for gray matter covariation between the CS and TTG. Second, we analyzed diffusion tensor imaging (DTI) from human connectome project (HCP) participants and conducted a voxel-based morphometry (VBM) meta-analysis to determine if white matter pathways connect the CS and TTG. Third, we analyzed 500 HCP participants to evaluate whether task-free functional connectivity between the CS and TTG. Fourth, we similarly examined whether task driven, natural audiovisual speech perception results in functional connectivity between the CS and TTG. Fifth, we used a coactivation meta-analysis to evaluate whether the CS and TTG tend to be coactivated in the same sets of studies and whether those studies are lower-level speech production and perception related. Finally, we conducted more exploratory analyses to determine if genes overexpressed in the hypothesized CS and TTG circuit are described as being involved in lower-level sensorimotor functions, vocal learning, and disorders of efference copy and prediction. In all analysis, we contrasted TTG connectivity with that of other nearby auditory regions (like the planum polare, planum temporale, and STG) to examine whether the CS-TTG circuit is separable from networks formed by adjacent regions.

## 2 Methods

### 2.1 Gray matter structural connectivity

We first evaluated whether the human CS and TTG show gray matter thickness covariation. Prior work examining cortical thickness correlations involving the lateral temporal cortex and superior temporal plane (STP) has typically focused on deriving whole-brain modular structure from functional connectivity maps with a low degree of resolution, e.g., treating the entire STG as one region (Z. J. Chen, He, Rosa-Neto, Germann, & Evans, 2008)(see Atilgan, Collignon, & Hasson, 2017 for recent exception). At this level, no connections between auditory and the primary motor/premotor regions have been reported. However, this could be due to averaging over multiple subregions of the STP, which have markedly different functionality in terms of bandpass characteristics (Hall et al., 2002), sensitivity to speech vs. nonspeech sounds (Leaver & Rauschecker, 2010), and role in coding the location vs. identity of auditory objects (see Bizley & Cohen, 2013). Thus, to evaluate the hypothesis that the CS and TTG are structurally connected, we used a finer granularity of STP parcellation, following a protocol we previously developed in which the STP is parcellated into 13 subregions using manual anatomical definitions (Tremblay et al., 2013)(Figure 1, left).

#### 2.1.1 Participants

Structural data used for this analysis were obtained from 33 volunteers who had participated in functional studies of normal auditory functions. All participants were interviewed by a board-certified medical doctor prior to participation. The interview screened for exclusion criteria including those based on physical and mental health. Data collection was approved by the Institutional Review Board (IRB) of The University of Trento.

#### 2.1.2 Imaging parameters

A 4T 8-coil Bruker system was used to acquire high-resolution anatomical and functional data for each participant. Structural scans were acquired with a 3D T1-weighted MPRAGE sequence (TR/TE = 2700/4 ms; Flip angle = 7°; Matrix = 256 × 224; 176 sagittal slices; Isotropic voxel size = 1 mm^3^). Two structural volumes were obtained for all participants, one optimized for signal-to-noise ratio and one for contrast-to-noise ratio.

#### 2.1.3 Imaging analyses

The two structural images were averaged to allow more accurate structural image processing via the FreeSurfer 5.3.0 pipeline (Fischl, 2012). They were evaluated by UH, who carried out manual interventions when the initial segmentation resulted in inaccurate white-matter/gray-matter segmentation (including control-point intervention to manually adjust distributions of white- and gray-matter intensity, manual addition of white matter, or manual removal of residual dura mater or skull). Whole-brain cortical thickness measures were obtained from FreeSurfer routines calculating the closest distance from the gray/white boundary to the gray/CSF boundary at each vertex on the tessellated surface. These individual maps were exported to SUMA (AFNI’s 3D cortical surface mapping module)(Cox, 1996; Saad, Reynolds, Argall, Japee, & Cox, 2004a). To maintain a common vertex numbering scheme for all participants, we created a new version of each participant's mesh using the *MapIcosahedron* utility. We then smoothed the individual-participant cortical thickness data on the surface using a heat-kernel smoother (8 mm FWHM). The purpose of this moderate smoothing was to improve the alignment of cortical thickness values across participants.

The cortical thickness values for the STP seed regions were obtained from the non-smoothed, original surfaces on which the 13 regions of interest (ROIs) were annotated per hemisphere using a previously defined parcellation (P. Tremblay, Baroni, & Hasson, 2013). For a given seed region, a single vector reflecting all participants’ mean cortical thickness in that region was constructed. Then, on a whole-brain level, this vector was correlated with similar vectors constructed for each single cortical *vertex*. We included SPT regions themselves in this analysis to evaluate the degree of correlation within the whole region. Correlation maps were computed using robust regression, implemented via the *R* package *robustbase* (Team, 2014; Todorov, Filzmoser, & Others, 2009). The seed-region value was the predicting variable, and each other vertex was the predicted variable. Control for family-wise error (FWE) was implemented via cluster-based correction (p < .005 single-voxel cluster forming threshold, p < .05 FWE based on simulations on the surface implementing a 10 mm inherent smoothing).

Prior to conducting the whole-brain analysis against seed-region data we evaluated whether there was dependence in the cortical thickness vectors across the 26 STP subregions (13 per hemisphere). If there were strong inter-regional correlations in this value, this would mean a single-region analysis is not licensed as whole-brain correlations would also be highly correlated. To evaluate this, we analyzed the 33 (subject) × 26 (region) value cortical thickness matrix using PCA, whose two component solution suggested weak interdependence among regions, as we report in the Results section.

### 2.2 White matter structural connectivity

To evaluate the extent to which human CS and TTG are connected by white matter pathways, we used a combination of analyses involving 1) tractography from a group of 40 adults from the HCP (Q3 data release) and 2) meta-analyses of tractography studies. Using these two approaches, we evaluated whether white matter connections from the CS and TTG overlap in one or both of these regions as they would if there is some connection between them (though perhaps mediated by multiple synapses). For comparison, we also evaluated the overlap of nearby premotor region connectivity and the connectivity of the TTG though they were not expected to be interconnected.

Specifically, HCP tractography data were processed as part of the publicly available “Human Brainnetome Atlas” (Fan et al., 2016). The atlas partitions the brain into 246 regions on the basis of tractography connectivity architecture. We created three binary masks from these data, based on probabilistic tractography results thresholded at .95 (where the range is 0-1). These consisted of a BA 41/42 mask (comprised of four regions: bilateral BA 41/42 and TE1.0/TE1.2 parcellations), BA 4 mask (10 regions, five bilateral BA 4 parcellations), and premotor BA 44/6 mask (16 regions, 5 bilateral BA6 parcellations and 3 bilateral BA 44 parcellations).

Separately, we conducted two meta-analyses of white matter connectivity from BA 4 and BA 41/42 using the BrainMap VBM database (http://brainmap.org/). Specifically, we queried the database (in October, 2016) for all experiments that met the “Experiment” tissue “Contrast” type of “White Matter” and that involved connectivity from “Locations” with a Talairach Daemon or “TD Label” that is from the “Cell Type” of either “Brodmann area 4” or “Brodmann area 41 or 42”. These queries returned X/Y/Z stereotaxic coordinate space “Locations”, that is, centers of mass of structural brain data reported in neuroimaging papers (Fox et al., 2005; Fox & Lancaster, 2002; Laird, Lancaster, & Fox, 2005). Locations that were originally published in the Talairach coordinate space were converted to Montreal Neurological Institute (MNI) space (Laird et al., 2010; Lancaster et al., 2007). We then edited each of the two resulting location files to keep only contrasts for “Normals” or “Normals” greater than some non-“Normals” group.

We then performed Activation Likelihood Estimation (ALE) meta-analyses for the BA 4 and BA 41/42 files by modelling each MNI location as a three-dimensional probability distribution and quantitatively assessing their convergence across experiments. Significance was assessed by permutation analysis of above-chance clustering between experiments (S. B. Eickhoff, Heim, Zilles, & Amunts, 2009; Simon B. Eickhoff et al., 2011; Simon B. Eickhoff, Bzdok, Laird, Kurth, & Fox, 2012; Turkeltaub et al., 2012). The conjunction (i.e., overlap) of the two ALE meta-analyses was created using ten thousand permutations to derive p-values (Simon B. Eickhoff et al., 2011). All resulting ALE maps were false discovery rate (FDR) corrected for multiple comparisons to a combined p < 0.05 and further protected by using a minimum cluster size of 800 mm^3^ (100 voxels).

Finally, we created white matter overlap maps for the HCP data and meta-analyses to visualize across region white-matter structural connectivity. In particular, we formed the union of the 1) BA 41/42 and BA 4 and 2) BA 41/42 and BA 44/6 HCP binary maps, and 3) BA 41/42 and BA 4 BrainMap meta-analyses binarized maps. We then combined the resulting overlap maps to visualize where overlap maps overlap from two different approaches with the expectation that the two BA 41/42 and BA 4 maps (but not the BA 41/42 and BA 44/6 map) should overlap in the CS and/or TTG.

### 2.3 Task-free functional connectivity

To address whether the CS and TTG form an intrinsic functional network while the brain is not engaged in an explicit task, we analyzed resting-state fMRI from the publicly available WU-Minn HCP “900 Subjects Data Release” (Marcus et al., 2011; Van Essen et al., 2013). All structural and functional MRI data had been preprocessed by the HCP using Version 3 of the HCP preprocessing pipeline as described below (Glasser et al., 2013).

#### 2.3.1 Participants

We pseudo-randomly selected 500 of the 900 participants. Each participant was required to have an anatomical parcellation, four resting state scans over two sessions, each with associated physiological (respiratory and cardiac) monitoring files.

#### 2.3.2 Procedure

Each of the four resting state scans was 14 minutes and 33 seconds long. Participants were instructed to keep their eyes open and fixate on a light cross-hair on a dark background in a darkened room. Head motion was monitored using an optical motion tracking camera system. Cardiac and respiratory signals were monitored using a pulse oximeter placed on a digit and a respiratory belt placed on the abdomen, sampled equally at 400 Hz.

#### 2.3.3 Neuroimaging

Participants were scanned on a Siemens 3T scanner using a standard 32-channel head coil.

##### Anatomical imaging and preprocessing

High resolution T1w 3D MPRAGE structural scans were acquired (TR = 2400 ms; TE = 2.14 ms; T1 = 1000; Flip angle = 8°; FOV = 224 × 224 mm; Voxel size = 0.7 mm isotropic). Preprocessing involved gradient distortion correction, coregistration and averaging of T1w and T2w runs, ACPC (i.e. 6 dof) registration for T1w and T2w, initial atlas based brain mask extraction of T1w and T2w, field map distortion correction and registration of T2w to T1w, bias field correction using sqrt (T1w X T2w)(Rilling et al., 2011), and nonlinear registration to the MNI152 atlas. A FreeSurfer pipeline (based on FreeSurfer 5.3.0-HCP)(Fischl, 2012) was then used to segment the volume into predefined structures, reconstruct white and pial cortical surfaces, and perform FreeSurfer’s standard folding-based surface registration to their surface atlas (fsaverage). All images were defaced before being made available (Milchenko & Marcus, 2013).

##### Functional imaging and preprocessing

In each session, oblique axial images were phase encoded in a right-to-left (RL) direction in one run and the left-to-right (LR) direction in the other. Gradient-echo EPI was used to collect 1200 images per run, acquired using multiband accelerated imaging (D. A. Feinberg et al., 2010; Moeller, Yacoub, & Olman, 2010; Setsompop et al., 2012; Xu, Moeller, Strupp, Auerbach, & Chen, 2012)(TR = 720 ms; TE = 33.1 ms; Flip angle = 52°; FOV 208 × 180 mm; 2.0 mm slice thickness; 72 slices; 2.0 mm isotropic voxels; Acceleration factor of 8).

Preprocessing of timeseries as completed by the HCP consortium included gradient distortion correction, motion correction using the “SBRef” volume as the target (Smith SM et al., submitted), TOPUP-based field map preprocessing using spin echo field map (for each day of each BOLD run), distortion correction and EPI to T1w registration of the “SBRef” volume, one step spline resampling from the original EPI frames to atlas space including all transforms, intensity normalization to mean of 10,000 and bias field removal, and masking using the FreeSurfer segmentation. Resulting timeseries were highpass filtered and cleaned of structured noise using the FSL independent component analysis (ICA) program MELODIC with FIX, a program trained on HCP data that automatically labels and removes artifactual components from resting state timeseries (Griffanti et al., 2014; Salimi-Khorshidi et al., 2014; Smith et al., 2013). In addition, 24 confound timeseries derived from the motion estimation (6 rigid-body parameter timeseries, their backwards-looking temporal derivatives, plus all 12 resulting regressors squared)(Satterthwaite et al., 2013), were used to remove nuisance effects from the timeseries. These motion parameters also had the temporal highpass filtering applied to them and were then regressed out of the data.

Finally, to evaluate the degree to which connectivity was driven by physiological factors, we created a set of 13 physiological regressors for each participant using *mcretrots* available in AFNI (Cox, 1996). Regressors included the respiration volume per unit time (RVT) and delayed terms (regressors 0-4), respiration (regressors 5-8), and cardiac (regressors 9-12)(Jo, Saad, Simmons, Milbury, & Cox, 2010).

##### Functional connectivity analyses

The analysis that we conducted began with the preprocessed timeseries. First, we created a set of 12 ROIs for each participant from that person’s Freesurfer anatomical parcellation. These included, bilaterally, the planum polare, TTG, transverse temporal sulcus (TTS), planum temporale, lateral fissure, and STG. Second, we averaged the timeseries from all voxels in each ROI to create a seed timeseries for each of the 12 ROIs, for each of the two phase encoding directions, and each of the two sessions. Third, we correlated each of the 12 seed time series with every other voxel in the brain for the phase encoding directions and sessions. We also conducted a separate correlation analysis using only the nuisance physiological regressors (this captured to what extent each voxel’s time series covaried with physiological fluctuations). Fourth, we z-transformed the correlation coefficient in each voxel for each of these maps. Fifth, we averaged the resulting maps over phase encoding direction and hemisphere and averaged together the TTG/TTS ROIs (for ease of presentation and likely inability to resolve differences between these regions). Thus, averaging resulted in in five maps per participant per session reflecting whole-brain functional connectivity of STP regions. Sixth, paired t-tests were run contrasting pairs of regions, using the physiological correlation maps as covariates for inter-individual differences to project the physiological related activity out of the data (see *3dttest++* in AFNI for further information). Finally, we also contrasted ROIs with the physiological correlations to assure effects were not in the CS-TTG regions. Given the large number of participants, we a priori chose a very high but ultimately arbitrary threshold of p = 1.00e-10 (corresponding to t = 6.61; df = 498) to threshold resulting groups maps and a minimum cluster size of 160 mm^3^ (or 20 voxels). To help assure that resulting differences were meaningful, we also arbitrarily required there be a minimum difference between contrasts of regions of 0.04.

### 2.4. Task-based functional connectivity during natural listening

To determine if the CS-TTG connectivity is manifest during natural audiovisual speech perception and language comprehension and not simply when the brain is at rest, we performed functional connectivity analysis using fMRI data collected while participants watched a television game show.

#### 2.4.1 Participants

There were 14 participants (6 females, 8 male; average age = 24.6, SD = 3.59 years). Each was a native English speaker, right-handed as determined by the Edinburgh Handedness Inventory (Oldfield, 1971) with normal or corrected to normal hearing and vision and no history of neurological or psychiatric illness. The study was approved by the Institutional Review Board (IRB) of Weill-Cornell Medical College and participants provided written informed consent.

#### 2.4.2 Stimulus and task

Participants listened to and watched 32 minutes and 24 seconds of an episode of a television game show (“Are You Smarter Than A 5th Grader”; Season 2, Episode 24, Aired 2/7/08). The episode was edited down from its original length without decreasing intelligibility. It was presented in six segments so that we could check on participants and give them breaks. Among other reasons, this show was chosen because it has a number of desirable properties including: Natural audiovisual dialogue between the host, contestant, and, to a lesser extent, six peripheral individuals and a built-in behavioural assay to assess participants knowledge of the items or events discussed in the video and to determine whether participants were attending. Participants who completed this assay following the experiment (n=10) were on average 98% accurate and 82% confident in their answers when asked the contestant’s answers to 11 questions during the show. The contestant on the show answered all questions correctly except the last. Participants were 80% accurate when asked the contestant’s answer to the final question after having seen the show despite that only 30% indicated that they had known the answer to the question before viewing the show (z(9)=2.2473; p < 0.02).

#### 2.4.3 Imaging parameters

Brain imaging was done on a 3 Tesla scanner (GE Medical Systems, Milwaukee, WI). A volumetric MPRAGE sequence was used to acquire anatomical images on which landmarks could be found and functional activation maps could be superimposed (FoV = 24; Sagittal Slices = 120; Voxel size = 1.5 × 0.9 × 0.9 mm). Functional imaging used an EPI sequence sensitive to BOLD contrast (TR = 1500 ms; TE = 30 ms; Flip angle = 75° deg; FoV = 220 mm; Base resolution = 64; Axial Slices: 25; Voxel size = 3.45 × 3.45 × 5 mm). There were six consecutive functional runs with durations, in minutes:seconds of 5:36, 5:45, 5:42, 5:42, 4:51, and 5:51. Each run began with a 10.5 second black screen that faded out to allow magnetization to reach a steady state and these images were discarded.

#### 2.4.4 Preprocessing

Unless otherwise noted, preprocessing was done with AFNI (Cox, 1996). Anatomical images were corrected for intensity non-uniformity, skull stripped (Iglesias, Liu, Thompson, & Tu, 2011), non-linearly registered to an MNI template, and inflated to surface based representations using Freesurfer software to create an anatomical parcellation for later use as ROIs (Fischl, 2012). Functional images from the six runs were spatially registered in 3D space by Fourier transformation. A correction was applied for slice timing differences, and spikes (signal intensities greater than 2.5 standard deviations from the mean) were interpolated to less extreme nearby values. We then corrected each run for head movement by registration to the mean of the middle functional run. These were then aligned to the MNI aligned anatomical images. We masked each run to remove voxels outside of the brain, linearly, quadratically and cubically detrended the time series, normalized them (by making the sum-of-squares equal to one), and concatenated them. The resulting timeseries were submitted to ICA to locate artifacts in the data (Beckmann & Smith, 2004). In particular, resulting components were automatically labelled as not-artifacts, possible artifacts, or artifacts using SOCK (Bhaganagarapu, Jackson, & Abbott, 2014). Each was manually reviewed for accuracy. There were an average of 237.29 components and an average of 177.79 artifactual components (74.93%) across participants. The independent component time course associated with each artifactual component was removed from the timeseries at each voxel using linear least squares regression. Finally, we blurred the resulting timeseries to a smoothness of 6 mm (Friedman, Glover, Krenz, Magnotta, & FIRST BIRN, 2006).

#### 2.4.5 Network analysis

We conducted a network analysis mostly following the task-free functional connectivity analysis done with the HCP data as described above. First, we created the same set of 12 ROIs for each participant from that participant’s Freesurfer anatomical parcellation (bilateral planum polare, TTG, TTS, planum temporale, lateral fissure, and STG). Second, we averaged the timeseries from all voxels in each ROI to create a seed timeseries for each of the 12 ROIs. Third, we correlated each of the 12 seed time series with every other voxel in the brain. Fourth, we z-transformed the correlation coefficient in each voxel for each of these maps. These 12 maps reflected whole-brain functional connectivity of the aforementioned STP regions for each participant. Finally, at the group level we used paired t-tests to contrast ROI based connectivity maps, averaging over TTG/TTS ROIs and averaging over hemispheres (resulting in five contrasts). Thresholds were determined for each pairwise contrast separately by first masking the data and then used the residuals from *3dttest++* and the “randomsign” flag to simulate 10,000 null 3D results in the mask, and then using ‘3dClustSim’ to generate cluster-thresholds at a corrected alpha level of .05 (Cox, Reynolds, & Taylor, 2016). This mask included the STP ROI regions as well as the pars opercularis (POp) of the inferior frontal gyrus (IFG), precentral gyrus and sulcus, paracentral gyrus and sulcus, CS, and postcentral gyrus and sulcus (Figure 4 white outline). We used a masking approach because we had hypotheses specific to these regions and because of reduced power for this study compared to other analysis in the manuscript.

#### 2.4.6 Functional Specificity

A final analysis assured that network results were functionally specific, i.e., related to speech perception and language comprehension. In particular, we tested the hypothesis that seed region timeseries would be more correlated with timepoints when audiovisual speech could be heard than times when faces could be observed without accompanying speech. To accomplish this, we manually coded the onset of every word in the TV show and convolved them with a standard model of the hemodynamic response function (HRF; namely the “Cox special” from the AFNI program *waver*). For a visually and auditorily similar comparison, we coded the onsets of every instance when faces were on screen but speech could not be heard though other noises like music, clapping, screaming, etc. were audible. We convolved these “faces no-speech” onsets with the same HRF model. The two resulting ideal waveform timeseries files, “audiovisual speech” and “faces no-speech”, were then correlated with the five ROI seed timeseries used in the network analysis for each participant. The resulting product-moment correlation coefficient values (r) were converted to z values using the Fisher z-transformation and contrasted across participants using a paired t-test.

### 2.5. Task-based functional connectivity meta-analysis

To address whether the CS-TTG form a network when participants are engaged by a wide variety of stimuli and tasks, we performed a meta-analytic joint activation analysis across a large number of neuroimaging studies. This analysis assumes that, if a brain region frequently coactivates with another brain region over many studies and statistical contrasts, than those regions can be considered to form a network. Our analysis proceeded in three stages: First, we used *coactivation-based clustering* to identify distinct functional subdivisions (clusters) within the superior temporal cortex itself, which is a step that in and of itself provides a novel approach for understanding the functional organization of the STP. We then used these obtained clusters within the STP to perform coactivation based meta-analysis on the whole-brain level. This analysis showed, for each STP subcluster, which brain regions tended to coactivate across version 0.6 of the Neurosynth database. This version contains 413,429 activation peaks or centers of mass from 11,406 studies and terms that appear at high frequency in the abstracts associated with those studies (Yarkoni, Poldrack, Nichols, Van Essen, & Wager, 2011)(https://github.com/neurosynth/). Analyses were based almost entirely on Python code provided by de la Vega and colleagues (2016)(available at https://github.com/adelavega/neurosynth-mfc/ along with Neurosynth https://github.com/neurosynth/neurosynth).

#### 2.5.1 Coactivation-based clustering

To find distinct functional domains in the STP without strong a priori anatomical or functional assumptions, we clustered voxels in the STP mask based on their coactivation with all other voxels in the brain, as coded in the Neurosynth database. Initially, “activation" in each voxel was represented as a binary vector of length 11,406 (i.e., the number of studies). We used a one or zero to code for whether the voxel was or was not within 10mm of an activation focus reported in a particular study. To reduce computational load, we then used principal components analysis (PCA) to reduce the whole-brain dimensionality to 100 components, producing a [# voxels × 100] matrix. Then, using this reduced description, we calculated the (Pearson correlation) “distance” between every voxel in the STP mask with each whole-brain voxels’ PCA component. K-means clustering was applied to the resulting matrix to group the STP voxels into six clusters. We chose this number because it matches the number of anatomical regions that compose the STP mask used for other analyses after averaging (and assuming left and right hemisphere regions have some functional homology). We also conducted k-means clustering using 12 and 26 regions. Though these demonstrated increasing specificity of activity patterns, they were not substantially different with respect to the hypotheses tested herein (i.e., all three showed independent CS-TTG networks). Thus, we use the k = 6 parcellation for ease of exposition (Figure 5, white outline). Note that this only constituted the initial step of the analysis, producing an automatic clustering of STP. The rest of the analysis is described next.

#### 2.5.2 Coactivation networks

For each of the 6 identified clusters we conducted a whole-brain task-based meta-analytic functional coactivation analysis. The principle here was to perform a formal contrast between studies that activated each of the six clusters as compared to studies that tended to activate the other clusters. The resulting statistical maps identify voxels throughout the brain that have a greater probability of coactivating with any particular STP cluster. A two-way chi-square (χ^2^) test was used to calculate p-values for each voxel between the sets of studies and we thresholded the resulting images using an FDR of q < 0.01.

#### 2.5.3 Functional profiles

After obtaining the coactivation networks, we generated qualitative profiles of the activity patterns in the clusters from which coactivation networks derived. This analysis speaks to the functional specificity of the hypothesized CS-TTG network for speech production and perception. This was done by determining which lexical terms from the published manuscripts indexed by Neurosynth best predicted activity in each of the six STP clusters across all 11,406 studies. That is, this analysis determines whether a classifier could predict if a study activated a specific cluster, given the terms mentioned in that study. A naive Bayes classifier was trained to discriminate six sets of high frequency terms associated with activation in each cluster versus a set of studies that did not produce activation in that cluster. Fourfold cross-validation was used for testing and the mean score was calculated across all folds as a summary measure of performance. Models were scored using both accuracy and area under the curve of the receiver operating characteristic (AUC-ROC). The log odds-ratio (LOR) - the probability that a term is present in active versus inactive studies - from the naive Bayes models from each cluster was used to generate the functional profiles. The LOR indicates whether a term is predictive of activation in a given cluster.

Given the theoretical questions asked here, we plot the functional terms with the highest LOR for a set of speech production, perception, and language comprehension terms and a set of motor related terms not involving movement of the articulators for comparison. In particular, the “Speech Production” category was comprised of the terms “vocal”, “production”, “overt”, “speech production”, and “articulatory”. *“*Speech Perception” terms are “speech sounds”, “pitch”, “phonetic”, “listening”, and “acoustic”. “Hand/Arm/Foot Movement” terms are “reaching”, “hand movements”, “foot”, “finger movements”, and “action observation". “Language Comprehension” terms are “syntactic”, “sentences”, “sentence”, “semantics”, and “language comprehension”. Note that the order in which these terms are written are the same order as presented in Figure 5 reading clockwise starting from the “S” in speech production. We elected to leave them out of the figure to increase readability.

### 2.6. Network-by-gene descriptions

We conducted exploratory analyses to determine if any genes overexpressesd in the CS-TTG networks that we have identified across analyses have functional properties consistent with our hypotheses. In particular, are the genes overexpressed in the CS-TTG network described in terms associated with lower-level sensorimotor processing, disorders associated with efference copy, and vocal learning? To address these questions, we developed a text mining approach to characterize the phenotype of sets of genes overexpressed in networks shown in Figures 3-5. The logic is that there exists a complex set of genes that regulate brain networks supporting language, though many studies focus on one gene or another (e.g., FOXP2). Rather than focus on specific genes, we assume that if researchers consistently describe specific genes in relation to particular speech and language related terms, and, furthermore, if those genes are particularly related to specific brain regions, than we can produce a functional profile of those regions reconstructed across a constellation of genes. This also provides a new picture of the sets of genes functioning in individual brain networks and their association with disorders of speech and language.

#### 2.6.1 Composite maps

To do this, we first created three summary brain images by combining the task-free and the two task-based functional connectivity analyses (Figures 6, top). From the resting state analysis, we used the maps as they appear in Figure 3 with areas in red assigned to a CS-TTG map and the rest into the Other map. For the television-viewing results, we used the maps as they appear in Figure 4 with hot colors assigned to the CS-TTG map and cool colors to the Other map. We excluded the lateral fissure contrast as it did not make contact with TTG despite having significant CS activity. Finally, for the meta-analytic connectivity data, clusters K6 and K3 as they appear in Figure 5 were assigned to the CS-TTG map and clusters K4 and K1 to the Other map. We excluded clusters K5 and K2 as they made less contact with TTG and because the functional analysis suggested they were less related to speech production and perception and more related to hand, arm, and foot movements. We scaled the data in each pair of (i.e., CS-TTG and Other) maps to range from 0-100 and combined them using the maximum value in each voxel (running analyses with the mean makes little difference). We clustered each map to minimum of 100 voxels. We divided the resulting map into three separate images: 1) Regions active only in the CS-TTG analyses; 2) A more inclusive CS-TTG map that include regions that overlapped with Other regions (the CS-TTG+); and 3) Regions active only in the Other map.

#### 2.6.2 Genes and descriptions

We formally cross-referenced these CS-TTG, CS-TTG+, and Other maps with gene expression data from six brains obtained from the Allen Human Brain Atlas (Hawrylycz et al., 2012). For every gene and each brain in the atlas, a linear model was fit to determine map similarity using gene expression decoding (Gorgolewski et al., 2014) as implemented in Neurovault (Gorgolewski et al., 2015). In particular, a one sample t-test was used to evaluate statistical significance using an FDR correction to account for the number of genes tested. We collated all significant genes, further correcting the FDR threshold for running the analysis on three brain images (.05/3; p < .01 FDR corrected).

We used a blind text-mining algorithm to find the functional descriptions of gene candidates overexpressed in these three networks. We conducted a PubMed search for each gene returning all titles, abstracts, and MeSH terms for that gene. We then made all resulting text lower case, removed all English stop words (e.g., “a” and “the”), and stemmed the document (e.g., “acoustics” and “acoustic” become “acoust”). We then counted the number of occurrences of each word and calculated the term frequency of that word per network (e.g., the term “cell” appeared 1,662,528 times in the Other network and was the most frequent term at 1.86% of words; compare this to the 596,468 appearances and term frequency of 1.78% in the CS-TTG network).

We then created a dictionary of 150 words related to five general categories: 1) Structural terms referring to body parts and brain regions linked to speech production and perception; 2) Motor terms directly referring to moving the body; 3) Sound and speech terms, mostly “lower-level”; 4) Terms specific to vocal learning, including reference to other vocal learning animals (e.g., songbirds); 5) Terms specific to language, mostly “higher level”; and 6) Terms associated with damage or disorders resulting in speech and language difficulties (see Supplementary Materials for a full list of stemmed words in this dictionary). We then subsetted our term frequency list by this dictionary and conducted pairwise χ^2^ tests for the frequency of each term in each network compared to the other networks. We report all terms and associated genes that showed a significant effect, using a Bonferroni correction for multiple comparisons as a cutoff (.05/(3*150) = p < 0.0001; Note the results do not change when using Fisher's exact test).

#### 2.6.3 Description validations

Finally, we provide evidence that the genes associated with descriptions that are significantly more frequent in one network over another are actually meaningfully related to those descriptions. To do this, we collated the list of genes associated with resulting vocal-learning-related terms as over-described in the CS-TTG networks. We took the intersection of those genes with a second set of genes that prior work identified as being uniquely associated with song learning birds relative to vocal non learners in the robust nucleus of the arcopallium (RA), most similar to the human CS (Pfenning et al., 2014). Specifically, Pfenning et al. (2014) found 40 genes that contributed to the shared specialized gene expression between the finch RA and human CS. They also found 55 genes that contributed to the convergent shared specialized expression between the RA analogs of multiple vocal learners (budgerigar, finch, and hummingbirds) and human laryngeal motor cortex (also in the CS) and adjacent somatosensory laryngeal cortex. We also collated a list of genes associated with the terms “schizophrenia” and “autism” that describe the CS-TTG networks and intersected the resulting genes with a list of 108 schizophrenia-associated genetic loci from the Schizophrenia Working Group of the Psychiatric Genomics Consortium (2014) and 107 genes associated with autism (De Rubeis et al., 2014). In all cases, if our text-mining based approach is accurate, there should be a high overlap with the genes associated with terms from specific networks and the genes specified in these other studies. It is worth noting that some genes could also appear in other networks but that, by definition, they are over-described as relating to vocal learning, autism, or schizophrenia and assumed, therefore, to be overexpressed in that network.

### 2.7. Presentation

Unless otherwise notes, all neuroimaging results are displayed on the MNI-aligned “Colin27” brain (Holmes et al., 1998). Surface representations of this brain were made using Freesurfer (Fischl, 2012) and displayed using SUMA (Saad, Reynolds, Argall, Japee, & Cox, 2004b). The Colin27 brain was automatically parcellated into anatomical regions of interest (Destrieux, Fischl, Dale, & Halgren, 2010) to serve as a guide to the location of results on surface representations.

## 3 Results

### 3.1. Gray matter structural connectivity

We evaluated whether the CS and TTG show gray matter cortical-thickness covariance, suggesting structural connectivity. To do so, we identified brain regions whose cortical thickness were correlated with the cortical thickness of left medial TTG. Figure 1 (left) shows the parcellation for the superior temporal plane in this study, and the results of the PCA analysis on regional cortical thickness values (13 regions per hemisphere) of the 33 participants. There was relatively limited shared variance across the subregions, suggesting regions likely have different whole-brain connectivity. This licensed analyses of single regions. As shown in Figure 1 (right), structural connectivity of left medial TTG covaried strongly with cortical thickness of the CS on the left, and in the dorsal CS on the right. In addition (not shown in Figure 1) there were strong correlations with cortical thickness of the right cuneus and lingual gyrus (primary visual cortex). These connections were specific to the medial TTG. For example, the immediately adjacent left medial TTS seed region covaried with the anterior insula bilaterally, and the left posterior superior temporal gyrus (see Figure S1 for more examples of cortical thickness specificity).

**Figure 1.**
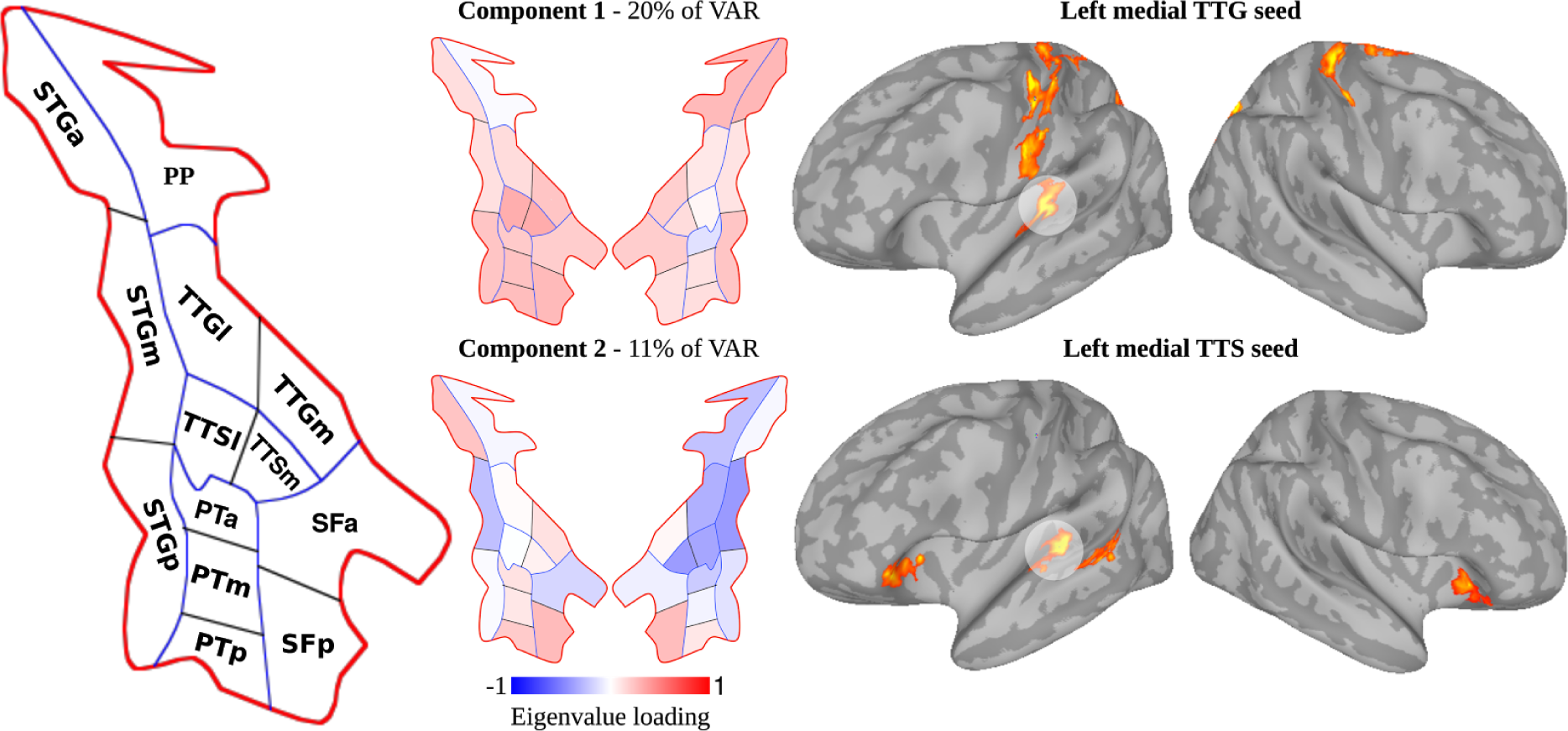
Design and analysis of structural connectivity analysis. We manually parcellated the superior temporal plane (schematic red outline) into 13 subregions. PP: Planum Polare, PT: Planum Temporale, STG: Superior temporal gyrus, SF: Sylvian Fissure. TTG: transverse temporal gyrus; TTS: transverse temporal sulcus. Subscripts: a/m/p = anterior/middle/posterior. m/l: medial/lateral. Mean cortical thickness values were derived per subregion, per participant, resulting in a 26 [regions] × 33 [participants] matrix. We evaluated the correlation between cortical thickness values in the 26 subregions of STP bilaterally using PCA. The PCA solution indicated there is no strong covariance amongst the regions. Demonstrating the specific cortical thickness covariance of medial TTG, strong correlations are found between left medial TTG and central sulcus bilaterally, whereas a different correlation structure is identified for the immediately adjacent left medial TTS. See Supplementary Materials Figure S1 for correlation maps for other left hemisphere STP subregions.

### 3.2. White matter structural connectivity

We evaluated whether the CS and TTG are connected by white matter tracts. If they are, the white matter pathways originating from BA 41/42 and BA 4 should overlap. We analyzed the overlap of B41/42 with immediately adjacent BA 44/6 for comparison using HCP DTI data and VBM meta-analyses. Figure 2 shows that both the BA 41/42 and BA 4 HCP (red) and meta-analysis (green) overlap maps resulted in connectivity around the medial aspect of the omega knob in the sagittal slices formed by the TTG. In addition, the two sets of maps themselves overlapped in this same region (yellow). In contrast, there was only a small overlap between the BA 41/42 and BA 44/6 overlap map and the other maps (teal). For the HCP data, overlap of connectivity in the frontal lobe was quite different for overlaps of BA 41/42 and BA 4 (red) and overlaps of BA 41/42 and BA 44/6 (light blue) and their overlap (darker blue). Specifically, only the BA 41/42 and BA 4 overlaps (red) covered the bulk of the TTG and made connections that terminated around the CS - compare coronal slice 61 in more frontal regions and slices 81 and 94 in more central regions (top row) or sagittal slices 65 and 67 (bottom row).

**Figure 2.**
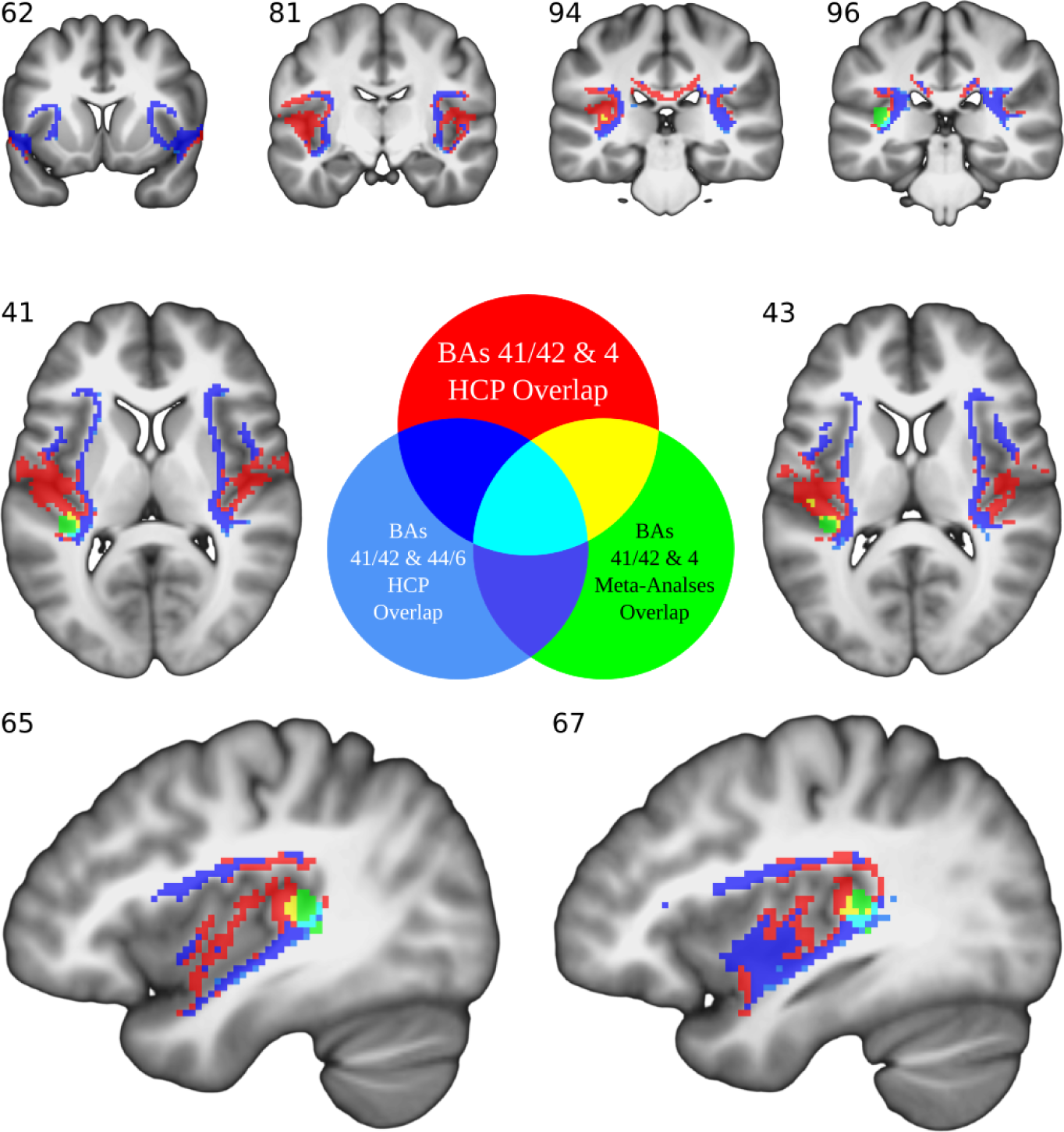
Overlap of white matter structural connectivity between cytoarchitectonic proxies for primary motor and primary auditory cortices. Overlap of Human Connectome Project (HCP) DTI data for BAs 41/42 and BA 4 (red) and BAs 41/42 and BAs 44/6 (dark blue). Overlap of Brainmap white matter meta-analyses for BAs 41/42 and BA 4 (green). The top row of coronal slices show that difference in connectivity are more anterior/posterior for BA 44/6 and BA 4 overlaps, respectively. The middle row of axial slices illustrates the significant overlap of BA 41/42 and 4 connectivity along the transverse temporal gyrus for the HCP data. Finally, the bottom row of sagittal slices shows significant overlap of BA 41/42 and 4 for both HCP and meta-analyses connectivity in the medial aspect of the left transverse temporal gyrus thought to correspond to primary auditory cortex.

### 3.3. Task-free functional connectivity

We evaluated whether the demonstrated CS and TTG structural connections might support “intrinsic” resting-state functional connectivity. Paired t-tests showed that task-free connectivity was significantly stronger between the TTG/TTS seed and CS than between the planum polare and CS bilaterally (Figure 3, red). The connectivity formed between the TTG/TTS and the CS was not significantly different from the connectivity formed between either the STG or planum temporale and the CS. Only the lateral fissure seed showed significantly more connectivity with the CS compared to TTG/TTS and this was mostly in the dorsal aspects (i.e., above a line extending the inferior frontal sulcus through the CS; Figure 3, black outlines). In contrast to these patterns, the STG seed formed significantly stronger connectivity with more anterior and posterior superior temporal regions and a small aspect of prefrontal cortex compared to the TTG (Figure 3, blue). The planum temporale formed significantly stronger connectivity with parietal cortices (Figure 3, green). These patterns of connectivity were nearly identical across sessions (compare Figure 3, Session 1 and Session 2).

To show that observed patterns are not a function of thresholding, we averaged over session and present the results at half the original p-value at decreasing levels of difference in correlations. These show patterns very similar to those already described except with an increasing pattern of overlap that, nonetheless, does not obscure TTG/TTS and CS interconnectivity (Figure 3, bottom). Furthermore, the greater STG connectivity with prefrontal cortices and IFG becomes more evident with decreasing r values (Figure 3, bottom, blue). Similarly, that the planum temporale forms more connections with the precentral gyrus and sulcus becomes more evident with decreasing r values (Figure 3, bottom, green). To determine if results are a physiological artifact related to prominent blood vessels in the CS, we ran paired t-tests using the results of the correlations run with the physiological files and ROI connectivity. All ROIs produced significantly stronger connectivity with the CS and sylvian fissure regions compared to physiological connectivity. Activity significantly greater for physiological files was mostly in the ventricles.

**Figure 3.**
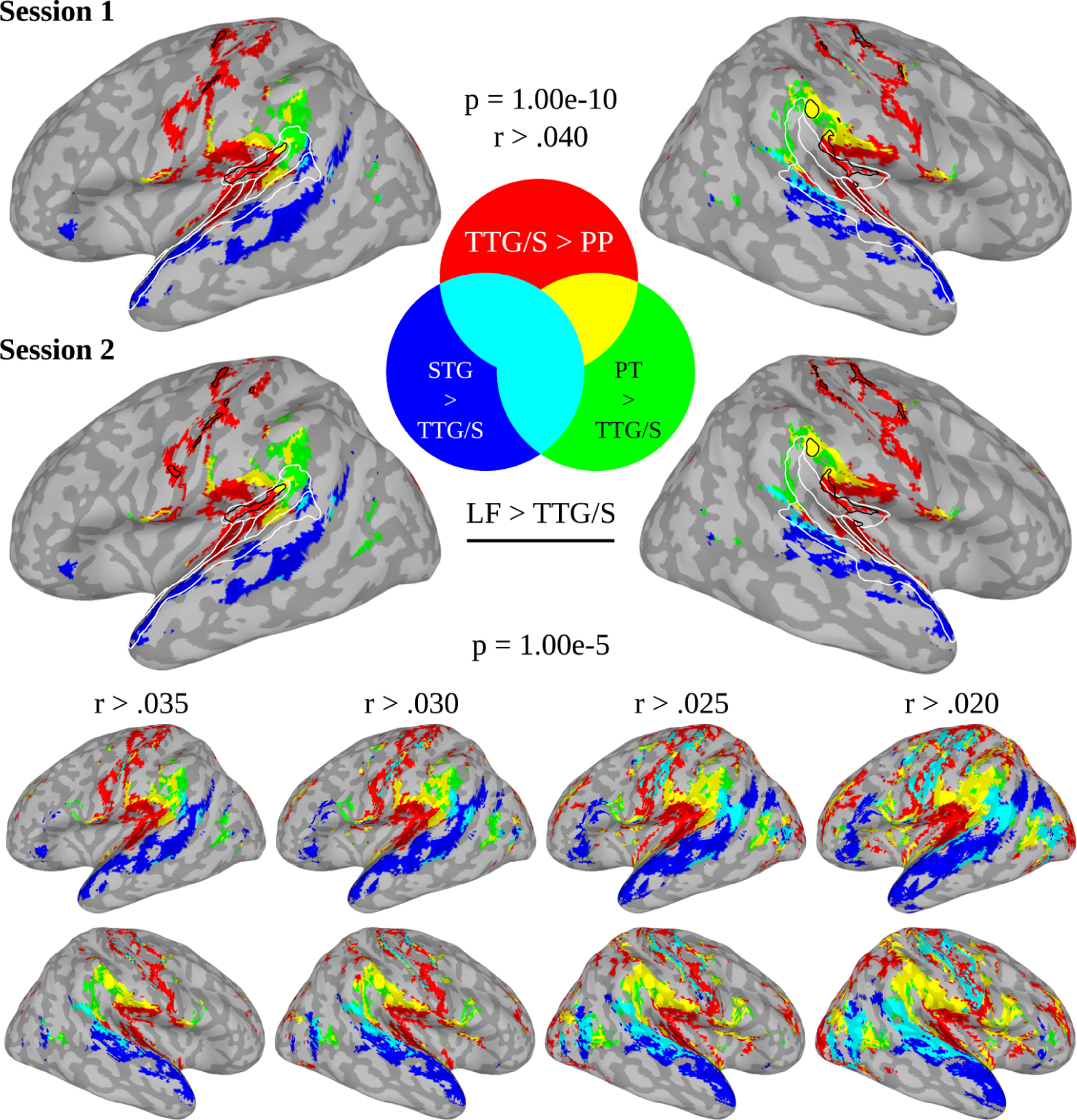
Task-free functional connectivity of the transverse temporal gyrus and sulcus (TTG/S) contrasted with other superior temporal seed regions. Analyses done with 500 participants’ Human Connectome Project “resting-state” data from two different days (Sessions 1 and 2). In both sessions and across multiple thresholds (bottom row), the TTG/S resulted in greater activity in the central sulcus (CS) than the planum polare (PP; red). In contrast, the TTG/S vs the superior temporal gyrus (STG; blue) or planum temporale (PT; green) produced more prefrontal/premotor activity. The only other contrast to produce CS activity was between the TTG/S and lateral fissure (LG; black outline) and this was mostly in dorsal CS.

### 3.4. Task-based functional connectivity during natural listening

We assessed whether the CS and TTG task-free connectivity extends to task-based functional connectivity associated with natural audiovisual speech perception. Figure 4 shows seed-based functional connectivity of the TTG/TTS while participants watched a television game show involving dialogue mostly between a host and contestants. The TTG/TTS resulted in significantly greater bilateral connectivity with the CS when compared to STG and planum polare connectivity (Figure 4, first and second rows, reds). The TTG/TTS also produced greater connectivity with the CS when compared to the planum temporale in the right hemisphere (Figure 4, third row, reds). In contrast to these patterns, the lateral fissure produced significantly stronger connectivity with the precentral sulcus and gyrus, CS, and postcentral gyrus and sulcus but not other STP regions (Figure 4, fourth row, blues). Furthermore, the STG and planum polare and temporale produced greater connectivity with more anterior and posterior superior temporal regions and the pars opercularis and precentral gyrus and sulcus in the frontal lobe, mostly bilaterally (Figure 4, rows 1-3, blues). As these analyses were done within a mask (Figure 4, white outline), we also present the unthresholded data to show how specific the TTG/TTS connectivity is with the CS and that results are not due to thresholding. Finally, connectivity patterns showed functional specificity for audiovisual speech. That is, the correlations between seed region timeseries and an ideal HRF waveform for “audiovisual speech” were significantly greater than “faces no-speech” (Table 1).

**Figure 4.**
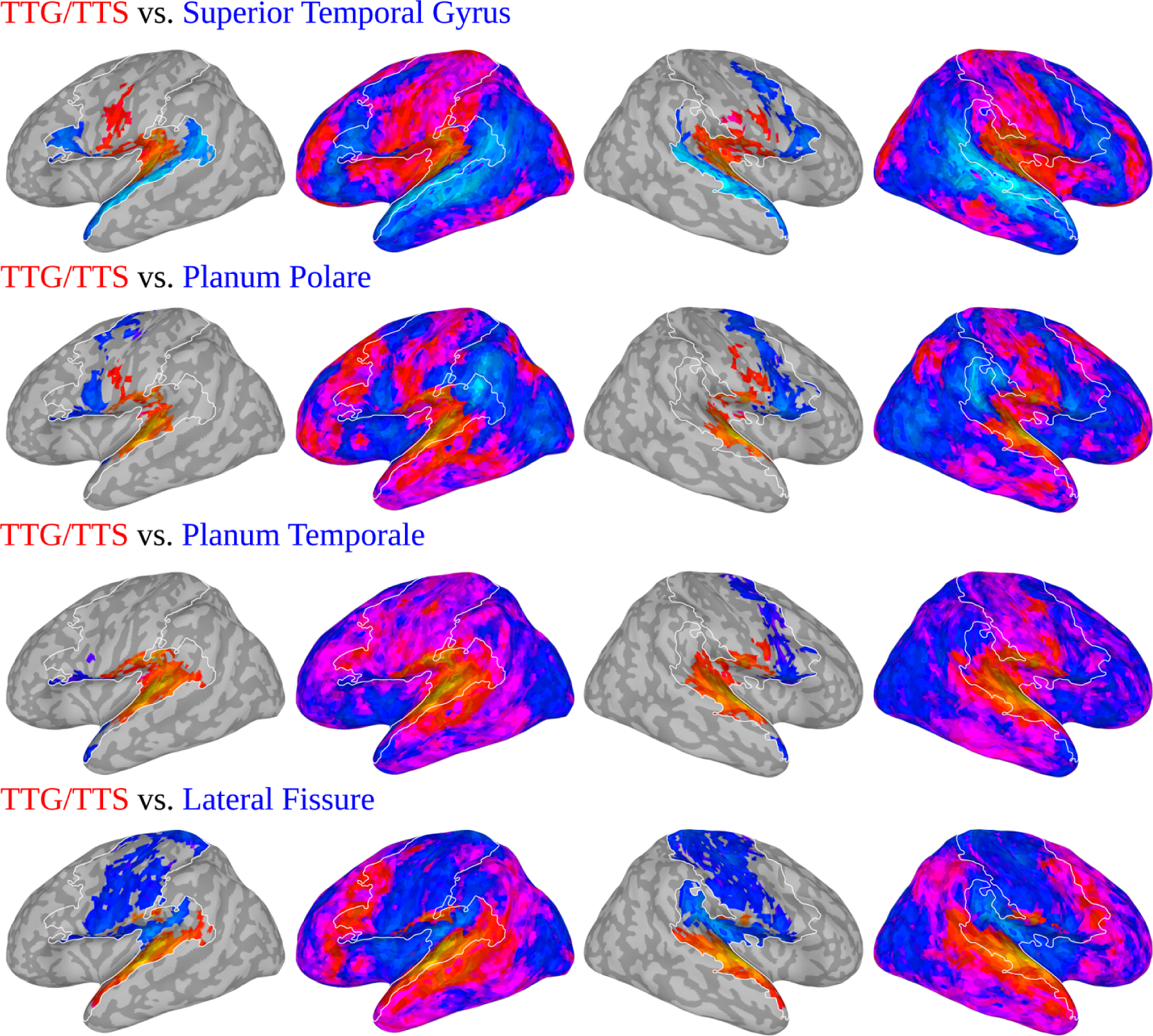
Functional connectivity of the transverse temporal gyrus and sulcus (TTG/S) contrasted with other superior temporal seed regions during television game show watching. Analyses presented both in a sensorimotor mask (white outline) at p < .05 corrected for multiple comparisons and unthresholded. With the exception of the lateral fissure, the TTG/S resulted in more activity in the central sulcus (CS; red) than all other regions (blues). Note that these results were speech specific. That is, superior temporal seed regions contributing to contrasts were significantly more correlated with audiovisual speech than time points in the show when faces and environmental noises were present (Table 1).

**Table 1.**
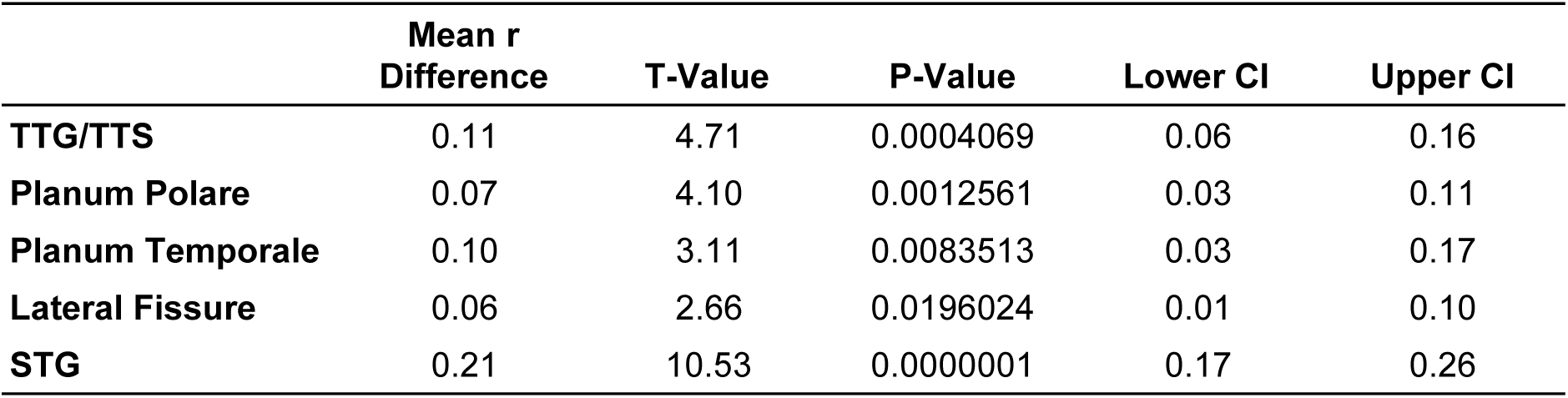
Contrast of the correlation between seed region timeseries with “audiovisual speech” and “faces no-speech” ideal hemodynamic response function waveforms.

### 3.5. Task-based functional connectivity meta-analysis

We next asked whether CS and TTG task-based functional connectivity extends to a large array task-based speech studies. The six clusters in the STP derived by unbiased coactivation-based clustering reasonably recapitulate standard parcellations of auditory cortex into primary and surrounding regions (Figure 5, left, white outlines). K-means clusters (K1-K6) included K1) the planum polare (-48/10/-17; 512 and 50/9/-19; 374 for MNI X/Y/Z centres of mass; numbers of voxels); K2) posterior lateral fissure and subcentral gyrus and sulcus (39/-24/15; 257 and -38/-28/14; 112); K3) transverse temporal sulcus and mid superior temporal gyrus (63/-14/-1; 480 and -61/-15/2; 297); K4) planum temporale and posterior STG (-59/-42/15; 367 and 63/-35/10; 173); K5) planum temporale and supramarginal gyrus (57/-38/26; 200 and -55/-40/26; 53); and K6) medial transverse temporal gyrus (51/-20/7; 220 and -47/-24/7; 213).

The whole-brain coactivation networks that were associated with these clusters were quite distinct (Figure 5, left). We used hierarchical clustering to organize them for display based on the similarity of their spatial patterns (Figure 5, far left). In this space, clusters K3 (lateral TTS) and K6 (medial TTG) were similar because they both coactivate with the ventral aspect of the CS (Figure 5, left, upper 2 rows). The planum temporale (K4) was mostly coactive with the pars opercularis and precentral gyrus and sulcus (Figure 5, left, third row). The planum polare (K1) was mostly coactive with the anterior IFG (Figure 5, left, fourth row). The inferior parietal cluster (K5) was coactive with mostly other parietal regions and precentral gyrus and sulcus (Figure 5, left, fifth row). Finally, the lateral fissure (K2) clusters was mostly coactive with parietal regions and the entire CS (Figure 5, left, sixth row).

Consistent with the different connectivity patterns, clusters K1-K6 had unique functional specificity profiles as evaluated by terms that discriminated coactivation patterns (Figure 5, right). Speech perception and production related terms predicted the activity in the TTG/TTS clusters (K6/K3) more than other terms (Figure 5, right, red lines). The terms resulting in the highest predictions were “pitch” and “speech sounds” in the “Speech Perception” category corresponding to the K6 cluster. “Vocal” and “speech production” were the highest in the “Speech Production” category for the K6/K3 clusters. In contrast, language comprehension related terms predicted activity in the planum temporale (K4) and planum polare (K1; Figure 5, right, green lines). Hand, arm, and foot related terms best predicted inferior parietal (K5) and lateral fissure (K2) activity (Figure 5, right, blue lines).

**Figure 5.**
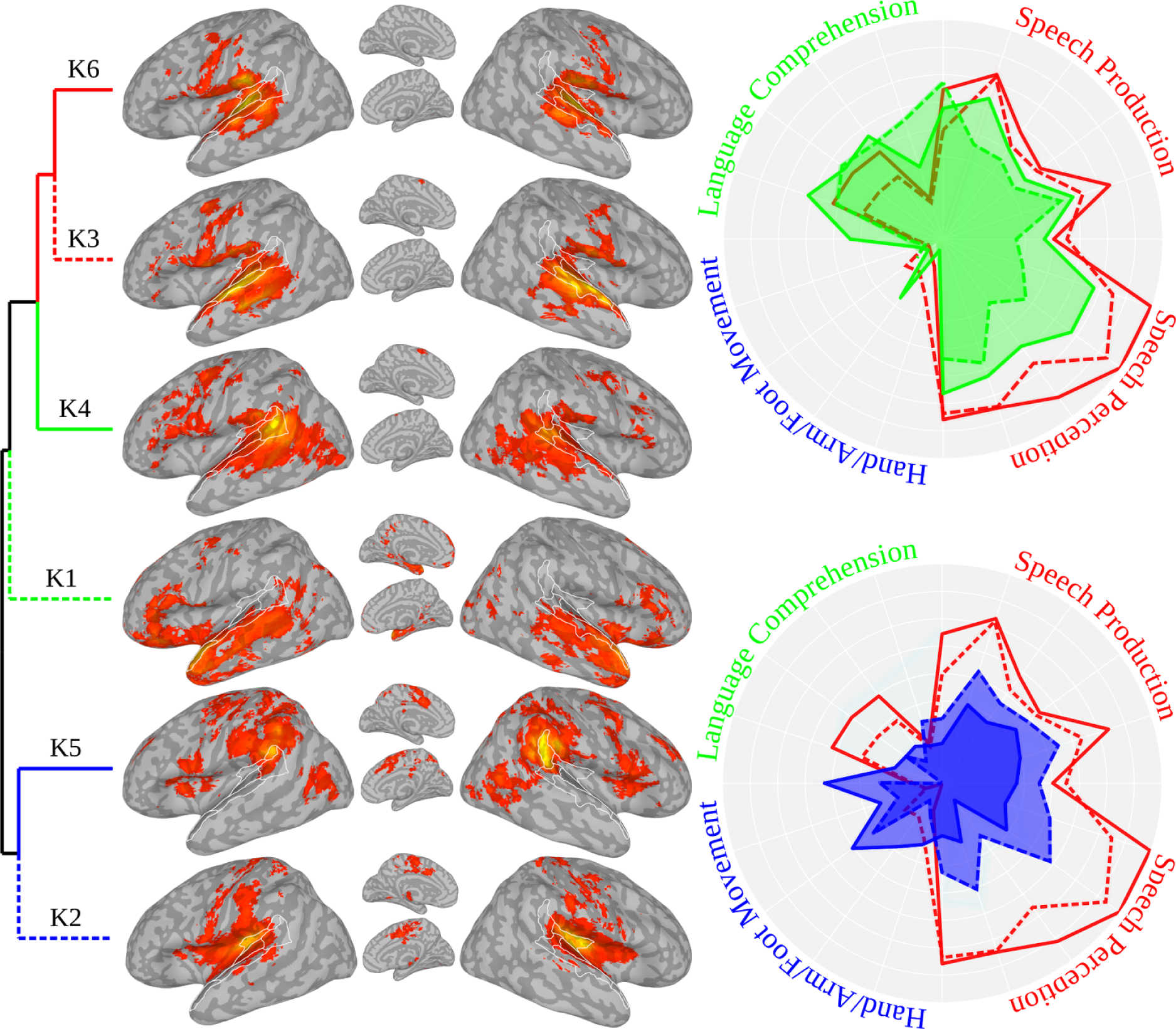
Task-based functional connectivity of superior temporal seed regions as determined my meta-analytic coactivation analysis. The superior temporal plane (white outline) was clustered into six regions based on coactivation patterns across 11,406 neuroimaging studies. Resulting clusters are presented in order of their coactivation similarly based on hierarchical clustering (left tree). In contrast to the other clusters, K6 and K3, originating from the TTG were coactive with the central sulcus. K6/K3 activity was more predicted by a set of speech production and perception terms than the other clusters (red). In contrast, clusters K4 and K1 were more associated with language comprehension terms (green) and K5 and K2 with movement of the distal limbs (blue). See Results Section 3.5 for MNI coordinate location of clusters.

### 3.6. Network-by-gene descriptions

Finally, we asked if the CS-TTG network overexpresses genes that have a speech and language functional phenotype more consistent with lower-level sensorimotor processing, vocal learning, and disorders connected with efference copy, as compared to other networks. To accomplish this, we first created summary network maps across analyses presented in Figures 3-5. The resulting CS-TTG map (i.e., network activity unique to TTG/TTS contrasts) is primarily located in the CS and the medial transverse temporal gyrus (Figure 6, black outline). The more permissive CS-TTG+ map (i.e., overlap of the CS-TTG and Other networks) includes these regions but also more of the superior temporal plane and dorsal CS (Figure 6, reds). The Other map (i.e., network activity unique to the planum polare, planum temporale, and STG contrasts) includes more posterior STG and middle temporal gyrus, anterior STG and temporal pole, inferior parietal lobule, precentral gyrus and sulcus, and inferior frontal gyrus regions (Figure 6, blues).

In total, the CS-TTG, CS-TTG+, and Other network maps were associated with 5,735 unique genes. The CS-TTG network was associated with 1,627 total and 357 unique genes. The CS-TTG+ and Other networks were associated with 2,997 (703 unique) and 4,475 (2,201 unique) genes, respectively. These sets of genes resulted in unique speech and language functional descriptions for each network as determined by a 150 term dictionary (see Supplementary Materials for dictionary and gene data). In particular, there were 55 terms that were significantly more frequent in one or more of the χ^2^ tests (Figure 6, word clouds). These break down into 44, 34, and 37 significant terms for the CS-TTG vs Other, the CS-TTG vs the CS-TTG+, and the CS-TTG+ vs Other χ^2^s, respectively.

These genes can be further broken down into sets of terms appearing more in one χ^2^ than appearing in other contrasts or the intersection of contrasts. There were 11 terms for which CS-TTG terms appeared at greater frequencies when contrasted with the CS-TTG+ and Other term frequencies *and* that were not more frequent in one of the converse contrasts. In pseudocode:

> *setdiff((“CS-TTG > CS-TTG+” & “CS-TTG > Other”), (“CS-TTG+ > CS-TTG” & “CS-TTG+ > Other” & “Other > CS-TTG” & “Other > CS-TTG+”))*

These were (where words in brackets provide further explanation):

> *audiolog*, *audiometri*, *binaur*, *charcot* [Charcot–Marie–Tooth disease], *genicul*, ***hvc***, *movement*, *otoacoust*, *syllab*, *tinnitu*, and *tonotop*

Using the same logic, there were 11 terms that were in the set of terms for which the CS-TTG+ terms appeared at greater frequencies. These were:

> *cleft*, *dysphagia*, *dyspraxia*, *grammat*, *languag*, *lip*, *palat*, *speech*, *stickler* [Stickler syndrome], *ttg*, and ***ultrason***

Finally, “*lexic*”, “*usher*” and “*william*” were the Other terms that appeared at higher frequencies.

> We also analysed at the intersection of the CS-TTG and CS-TTG+. In pseudocode: *intersect((“CS-TTG > CS-TTG+” & “CS-TTG > Other”), (“CS-TTG+ > CS-TTG” & “CS-TTG+ > Other”))*

This resulted in 22 terms:

> *arcuat,* ***autism***, ***bird****, cochlea, cochlear, cranial, deaf, ear,* ***gaba****, hear,* ***learn****, motor, muscl, nois,* ***schizophrenia***, ***song***, ***songbird****, teeth, tone, tooth,* ***vocal***, and *waardenburg* [Waardenburg syndrome]

Only “*ataxia*” was at the intersection of the CS-TTG and Other terms. Seven terms, “*angelman*” [Angelman syndrome] “*aphasia*”, “*oral*”, “*repeat*”, “*semant*”, “*somat*”, and “*word*”, were at the intersection of CS-TTG+ and Other.

**Figure 6.**
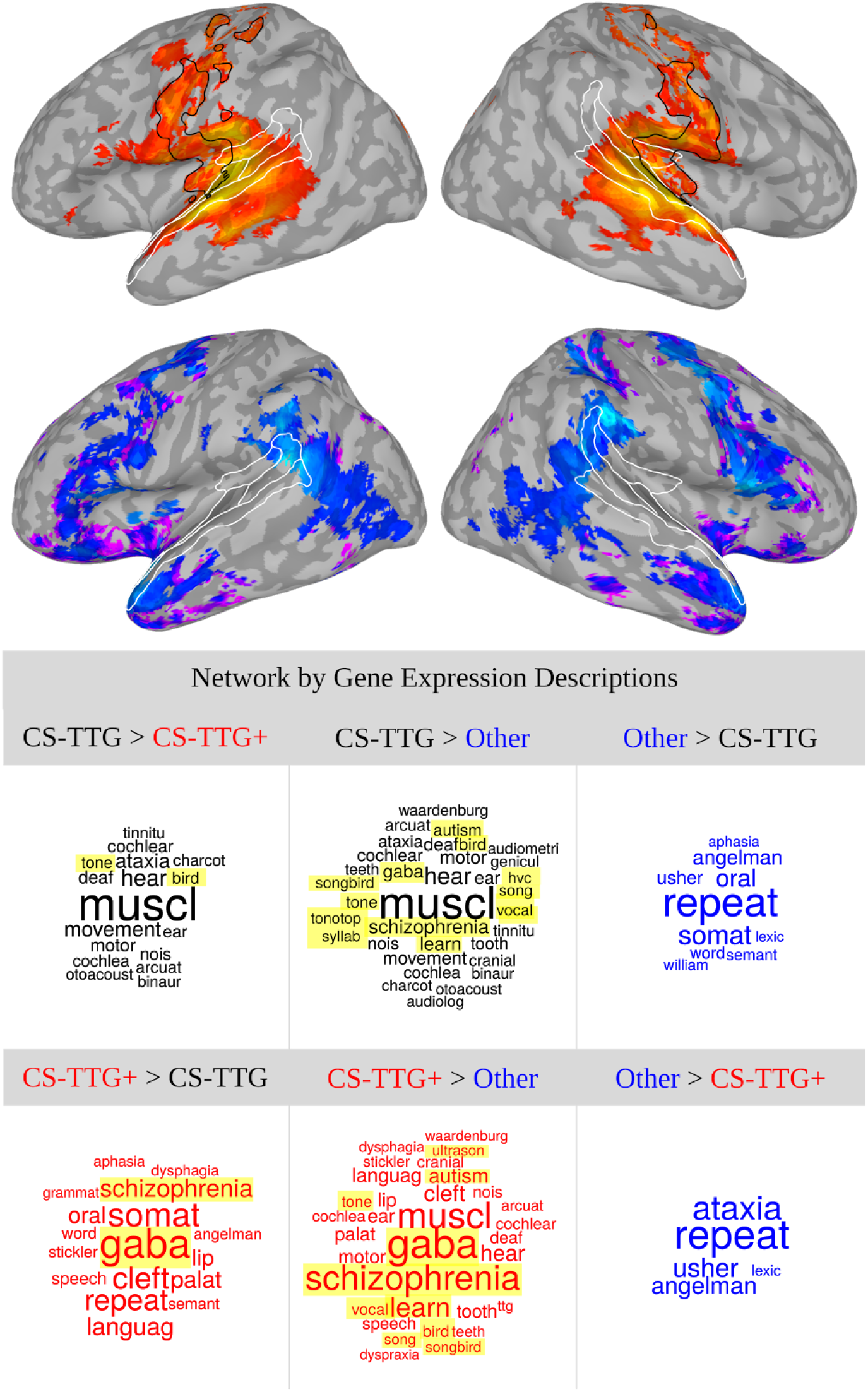
Composite networks across functional connectivity analyses (top) and gene based descriptions of those networks (bottom). Top. Composite networks were created by combining connectivity patterns from Figures 3-5 statistical contrasts. The resulting central sulcus and transverse temporal gyrus (“CS-TTG”) network encompasses connectivity patterns uniquely associated with the TTG/S (black outline). The “Other” network encompasses connectivity unique to the planum polare, planum temporale, and STG. Finally, the “CS-TTG+” network includes regions uniquely associated with the TTG/S contrasts and connectivity patterns overlapping with the “Other” networks. The white outline shows the location of superior temporal plane regions. Bottom. Word clouds are high frequency words from a dictionary of 150 speech-related terms (See Supplementary Materials). Specifically, each word cloud shows significantly more frequent term usage for the composite network for the contrast indicated. The CS-TTG and CS-TTG+ networks were more associated with low-level speech representations, vocal learning, and disorders associated with prediction and efference copy compared to the “Other” networks (terms highlighted in yellow).

Terms in bold for the CS-TTG and CS-TTG+ networks derive from an array of 1271 genes. These include those often associated with vocal learning, e.g., FOXP1 and FOXP2. To provide further evidence that these genes are associated with vocal learning and not simply a random assortment, we took the intersection of our 1271 genes with those documented by Pfenning et al. (2014) as uniquely associated with song learning bird and human CS/RA relative to non-learners. We found a 40% overlap with the genes specialized for song-learning birds and humans. Note that the chances of this happening randomly are *exceedingly* small. The probability of finding *only* nine genes while sampling 95 genes from 1271 would be about .0000005. It is even *less* probable when considering the total number of candidate genes we began with (5,735). Genes intersecting the 40 that contributed to the shared specialized gene expression between the finch RA and human CS where:

> *CNTN3, EPHA3, ERC2, GAP43, GDA, GRM5, HPCAL4, MARCH-1, MEF2C, MPPED1, NELL1, NELL2, PAK6, RAB26, SNCA, ZIC1*

Genes intersecting the 55 that contributed to the convergent shared specialized expression between the RA analogs of multiple vocal learners (budgerigar, finch, and hummingbirds) and human laryngeal motor cortex (in the CS) and adjacent somatosensory laryngeal cortex were:

> *B3GAT1, BCL11A, CHD5, CNTN3, DOCK4, GABBR2, GABRB3, GAP43, GPM6A, LINGO1, LYPD1, NECAB2, NEUROD6, PRPH2, SLIT1, SNCA, UCHL1, VIP, YWHAH*

We also collated a list of genes associated with the terms “schizophrenia” and “autism” (highlighted in red above) that describe the CS-TTG and the CS-TTG+ networks at a high frequency. We intersected the resulting 938 (for “schizophrenia”) and 700 (for “autism”) genes with a list of 108 schizophrenia-associated genetic loci (Schizophrenia Working Group of the Psychiatric Genomics Consortium, 2014) and 107 genes associated with autism (De Rubeis et al., 2014). The 35 intersecting schizophrenia genes where:

> ***AKT3****, ATP2A2, BCL11B,* ***CACNB2****, CD14,* ***CHRM4****, CHRNA3,* ***CSMD1****, CUL3, CYP26B1, DGKI,* ***DGKZ****, DPP4, EGR1, EPC2, FXR1, GFRA3, GPM6A, GRIN2A, HCN1, HSPD1, KCNB1, L3MBTL2, LRP1,* ***MAD1L1****, MEF2C,* ***MMP16***, ***NRGN***, ***PITPNM2****, RAI1, SATB2,* ***SBNO1****, SLC32TTG,* ***TCF4****, ZDHHC5*

The genes in bold are noted by the authors as having been previously associated with schizophrenia. The 17 intersecting autism genes where:

> ***ASXL3***, ***BCL11A****, CACNTTGD,* ***CUL3***, ***GABRB3***, ***GRIN2B***, ***KATNAL2****, KIRREL3,* ***MLL3****, MYT1L,* ***SCN2A****, SETBP1, SPARCL1, STXBP5,* ***TBR1****, TRIO, WHSC1*

Of these, 9 (in bold) where in the De Rubeis’ et al. (2014) top 22 most significant genes, themselves already known ASD risk genes. Again, the probability that we would find these sets of genes by chance is exceedingly small indicating the reliability of the network-by-gene description analyses.

## 4 Discussion

### 4.1. Summary

In six independent studies we present novel, consistent, and converging evidence for a core network of structural and functional connectivity between the central sulcus (CS) or primary motor and somatosensory cortex and the transverse temporal gyrus (TTG) or primary auditory cortex (Figure 6, top, black outline and red). Structural CS-TTG connectivity was verified by gray matter covariance and white matter connectivity in separate experiments (Figures 1 and 2, reds) and meta-analyses (Figure 2, green and yellow). These structural relationships had functional relevance in that the CS and TTG were functionally connected as shown in both task-free and task-based functional connectivity, again, in both experiments (Figures 3 and 4, reds) and large-scale meta-analyses (Figure 5, left, K6/3, reds). In all analyses, whether structural or functional, regions immediately adjacent to the TTG produced stronger connectivity with premotor and prefrontal regions. The general pattern was one in which more posterior superior temporal regions were more strongly associated with premotor connectivity whereas more anterior superior temporal regions produced more inferior frontal connectivity (Figures 1-6, blues).

Both of the task-based TTG connectivity analyses suggest that the CS-TTG circuit is strongly engaged by speech. First, both analyses showed greater ventral CS involvement compared to structural and task-free analyses, suggesting a greater engagement of motor areas that specifically innervate the articulators during speech related tasks (e.g., Conant, Bouchard, & Chang, 2014). Second, we found that the CS-TTG circuit is involved in real-world audiovisual speech perception. It showed greater responsiveness to words being spoken by visible talkers during a TV show than when faces were visible on screen and accompanied by natural environmental sounds (Figure 4, reds; Table 1). Finally, network meta-analysis suggests a general role for the CS-TTG network in both speech production and perception, with the latter particularly associated with “lower-level” speech sounds and pitch (Figure 5, radial plots, red). In contrast, meta-analyses of nearby and immediately adjacent superior temporal regions were more associated with moving the hands, arms, and feet or language comprehension (Figure 5, radial plots, blue and green).

Support for functional specificity was independently provided by network-by-gene description analyses (Figure 6, bottom, black and red). The genes expressed in the CS-TTG network were associated with “lower-level” sensorimotor terms at a higher frequency than they were for the other networks we defined. These include structural terms (like “arcuate”, “cochlea”, “muscle”, “palate”, and “ttg”), motor terms associated with body movement (like “motor” and “movement”), and the “lower-level” sound and speech terms “tone”, “tonotopy”, and “syllable”. The most prominent over-descriptions, however, were terms referring to vocal learning and vocal learning animals and, related, disorders sometimes associated with prediction and efference copy (Figure 6, bottom, yellow). As we review below, these included “learn”, “vocal”, “ultrasonic”, “bird”, “song”, “songbird”, and “HVc” (a songbird brain region) and “GABA”, “schizophrenia”, and “autism”. In contrast to these, genes expressed in networks associated with nearby and even immediately adjacent regions were described in terms of higher-level aspects of language like “lexical”, “semantic”, and “words” (Figure 6, bottom, blue).

These findings collectively suggest the pheno- and genotypic importance of this largely undocumented CS-TTG circuit. That is, the structural and functional connectivity, functional specificity, and related network-by-gene descriptions are consistent with the idea that this network supports vocal learning through efference copies that serve to rapidly activate “lower-level” acoustic targets for speech production and perception. In the case of speech production, we suggest the CS-TTG circuit supports vocal learning, speech target maintenance, and permits real-time adjustments to perturbations in production. In the case of speech perception, efference copy can constrain variability of the acoustic patterns arriving in auditory cortex, enabling perceptual constancy. In what follows, we present supporting evidence that CS-TTG structural connections might be new to humans and that functional connectivity associated with this new circuit supports activation of lower level representations through efference copy. We conclude by discussing implications for contemporary models of the neurobiology of speech perception and language comprehension that do not include this core circuit.

### 4.2. Structural connectivity

A rapid vocal learning, speech production and perception mechanism predicated on efference copy between the CS and TTG requires a structural connection mediated by few synapses. The structural analyses we conducted suggest that these connections exist. However, human *in vivo* structural connectivity measures are crude (Thomas et al., 2014), and the question remains whether there is evidence for direct primary motor and auditory connections as demonstrated by more precise antero- and retrograde tract-tracing methods. Prior work in several other animals suggests that, if these connections exist, they are sparse and that vocal learning, production, and perception processes largely rely on secondary or association cortices. For instance, in marmosets, a vocal learning primate (Takahashi et al., 2015), there is evidence for sparse projections between primary motor cortex and auditory association region CM but none with A1 (Burman, Bakola, Richardson, Reser, & Rosa, 2014). Furthermore, primary motor to auditory connections do not seem to be present in less vocal primates (e.g., Pandya, Gold, & Berger, 1969; Pandya, Hallett, & Kmukherjee, 1969). Mice are also moderate vocal learning animals albeit in the ultrasonic range (Arriaga, Zhou, & Jarvis, 2012), and they do have direct projections from primary and secondary motor cortices to auditory cortex (Nelson et al., 2013; Schneider et al., 2014). Still, when examined functionally with regard to efference copy, it is only secondary (M2) to auditory cortex connections that are described (Schneider & Mooney, 2015; Schneider et al., 2014).

For songbirds and parrots (arguably closest to humans in vocal learning skills)(Petkov & Jarvis, 2012), the available evidence also suggest limited connectivity between primary motor and auditory cortex. Instead, connectivity between the songbird homologue of primary motor cortex (the robust nucleus of the arcopallium, RA) and primary auditory cortex (field L) appears to be mediated by secondary auditory cortices (see Bolhuis, Okanoya, & Scharff, 2010) or other regions (Lewandowski, Vyssotski, Hahnloser, & Schmidt, 2013). While we have focused here on connectivity of primary motor cortex, the literature reveals similar patterns for primary somatosensory cortex (itself intimately connected with primary motor cortex)(Young, 1993). For example, in a recent review, Meredith and Lomber (2017) reported no cases of connectivity between primary somatosensory and auditory cortex in the marmoset and only sparse connections in rodents.

This relative lack of primary motor and somatosensory cortex and primary auditory cortex structural connectivity is in stark contrast to the human evidence that we present in Figures 1-2. However, some supporting human evidence is found in prior reports. Ro, Ellmore, and Beauchamp (2013) used DTI and showed direct connectivity in humans between primary auditory cortex and primary somatosensory cortex. Eickhoff et al. (2010) showed both structural and functional connections between primary motor cortex and operculum parietal (OP) 4, and functionally linked OP 4 to more basic sensorimotor as opposed to somatosensory processing. Finally, Sepulcre (2015) showed direct functional resting-state interconnectivity of primary motor and auditory cortices through OP 4. Sepulcre writes that his findings “introduce a fascinating possibility… of more straightforward communications between sensory-related and primary motor regions, without mediation by the classically defined dorsal and ventral language streams”. The data here strongly support the existence of this pathway and, in addition to connectivity mediated by OP 4, the possibility of direct primary motor or somatosensory connections with primary auditory cortex. Most importantly, as we summarized above, this pathway is also functionally relevant to speech.

### 4.3. Functional connectivity

Our structural connectivity data suggests that another organisational change in humans that make us so adept with speech, in addition to changes in laryngeal motor cortex connectivity reviewed in the Introduction, is a large increase in the CS-TTG structural connectivity. We proposed that the specific functional relevance of the CS-TTG structural connection is that it helps support vocal learning, speech production, and perception by providing a pathway for efference copy. Supporting a role in vocal learning are the network-by-gene descriptions which identified the terms “vocal” and “learn” as more strongly linked with the CS-TTG network than other networks. Other overexpressed terms included those associated with vocal learning animals like “ultrasonic” for mice (Arriaga et al., 2012), and “bird”, “song”, and “songbird”, associated with the prevailing nonhuman animal vocal learners (Petkov & Jarvis, 2012). Related to these, the CS-TTG network was linked to genes disproportionately associated with the term “HVc”, a brain region important for song learning in songbirds (Day, Kinnischtzke, Adam, & Nick, 2008; Lovell, Clayton, Replogle, & Mello, 2008; Mooney & Prather, 2005; Troyer & Doupe, 2000). Consistent with this vocal learning interpretation are the genes overexpressed in the CS-TTG network associated with these terms. That is, we documented a 40% overlap between a set of genes associated with the CS-TTG circuit on the one hand, and a second set of genes identified by Pfenning et al., (2014) as distinguishing song-learning birds and humans from non-learning birds (like SLIT1). This suggests that the genes overexpressed in the CS-TTG circuit are related to core vocal-related processes shared across phylogeny. Further supporting the vocal learning interpretation, we found that FOXP2 was overexpressed in the CS-TTG network. FOXP2 is thought to be required for vocal learning in both humans (Lai, Fisher, Hurst, Vargha-Khadem, & Monaco, 2001) and songbirds (Haesler et al., 2004, 2007; Scharff & Petri, 2011; Teramitsu, Kudo, London, Geschwind, & White, 2004).

Supporting a role for an efference copy mechanism in vocal learning are the network-by-gene descriptions that identified the terms “GABA”, “schizophrenia”, and “autism” as more strongly linked with the CS-TTG network than other networks. GABA, or γ-aminobutyric acid, is an inhibitory neurotransmitter associated with both vocal learning (Heaney & Kinney, 2016/4; Luo & Perkel, 1999; Vallentin, Kosche, Lipkind, & Long, 2016) and inhibitory aspects of efference copy (Bell & Grant, 1989; Mooney & Prather, 2005; Poulet & Hedwig, 2002, 2003, 2006). “Schizophrenia” is itself connected with GABA (Lewis, Hashimoto, & Volk, 2005; Stan & Lewis, 2012) and auditory hallucinations associated with schizophrenia are often claimed to be a disorder of efference copy (I. Feinberg, 2011; J. M. Ford et al., 2001, 2013; Judith M. Ford & Mathalon, 2005, 2012; Heinks-Maldonado et al., 2007; Pynn & DeSouza, 2013; Whitford, Ford, Mathalon, Kubicki, & Shenton, 2010). Indeed, several neuroimaging meta-analyses show primary auditory cortex is engaged during auditory verbal hallucinations (Kompus, Westerhausen, & Hugdahl, 2011) as is the somatosensory/postcentral gyrus, often overlapping with motor cortex (Kompus et al., 2011; Kühn & Gallinat, 2010; van Lutterveld, Diederen, Koops, Begemann, & Sommer, 2013; Zmigrod, Garrison, Carr, & Simons, 2016). Finally, the CS-TTG network was over-described by “autism”, a disorder, like schizophrenia, that is sometimes described as involving a failure of prediction, closely related to an efference copy mechanism (Friston, 2016; Pellicano & Burr, 2012; Rubenstein & Merzenich, 2003; Sinha et al., 2014). Supporting these interpretations, we found a high overlap of the genes associated with the terms “schizophrenia” and “autism” and those identified in other studies as schizophrenia and autism associated genes (De Rubeis et al., 2014; Schizophrenia Working Group of the Psychiatric Genomics Consortium, 2014).

Our findings are also consistent with the perspective that the sensory consequences activated by efference copy in the CS-TTG circuit are relatively low-level. Indeed, the functional connectivity meta-analyses suggest that the peak terms related to the CS-TTG network were “pitch” and “speech sounds” compared to language comprehension terms for the other networks. The network-by-gene analyses identified similar textual descriptions that were linked to genes diagnostic of the CS-TTG circuit. That is, we found that the CS-TTG circuit was more strongly related to genes associated with the terms “tone”, “tonotop”, and “syllab”, whereas the other networks we defined were, again, more associated with higher-level language terms like “lexical” “semantic”, and “word”. Thus, in both sets of analyses, terms are consistent with the finding that primary motor cortex is involved in pitch, rhythm, or ‘‘melodicity” across species during both production and perception (Arriaga et al., 2012; Brown et al., 2009; J. L. Chen, Penhune, & Zatorre, 2008; Merrill et al., 2012).

### 4.4. Implications

What are the implications of our current findings, taken together with the supportive recent work we reviewed, for current models of speech perception and language comprehension? We have shown that the CS-TTG circuit is realized via multiple types of structural connectivity, functional connectivity as well as evidence from coactivation meta-analyses. This in itself establishes the importance of understanding the computations carried out by this circuit. Based on the specificity of this connectivity pattern (and its divergence from contiguous regions like the planum polare, planum temporale, and STG), we argue that CS-TTG connectivity constitutes a “speech core” that could support low-level efference copy. This network is missing from contemporary “dual-stream” models of speech perception and language comprehension (Hickok & Poeppel, 2007; Rauschecker & Scott, 2009) though is partially represented in some models of speech production (Guenther et al., 2006; Hickok, 2012a). Thus, at minimum, this network will need to be integrated into some existing models.

We, however, suggest a different theoretical perspective that could account for the data. In particular, the CS-TTG circuit might develop relatively early and scaffold the development of higher-order speech comprehension systems. This is similar to more general models in which it has been argued that early to mature regions act as anchors or tethers that allow the topography of the rest of the brain to form through activity-dependent self-organization (Bourne & Rosa, 2006; Buckner & Krienen, 2013; Guillery, 2005; Rosa & Tweedale, 2005). Indeed, developmentally, primary motor and auditory cortex form earlier than other regions (Gogtay et al., 2004; Huttenlocher & Dabholkar, 1997; Qiu, Mori, & Miller, 2015). The CS-TTG circuit may be an anchor that is not only structural but also has the proper functional elements needed for learning thanks to seemingly expanded connectivity between the CS and TTG that permit efference copy as reviewed.

What does the topology of these other learning and activity-dependent self-organizing language networks look like? We suggest that CS-TTG speech core forms part of a core-periphery type organization like those recently used to describe complex social and brain networks, including language networks (Bassett et al., 2013; Borgatti & Everett, 2000; Chai, Mattar, Blank, Fedorenko, & Bassett, 2016; Csermely, London, Wu, & Uzzi, 2013). Core-periphery networks can account for the dynamic reconfiguration of functional modules with experience (Bassett et al., 2013). They can, therefore, account for a great deal of data demonstrating that the organization of (peripheral) brain networks are not fully stable, but dynamically reorganize with tasks and experience (Alavash, Thiel, & Gießing, 2016; Bassett et al., 2011, 2013; Bola & Sabel, 2015; Grandjean et al., 2017; Jeremy I. Skipper et al., 2017; Z. Wang et al., 2012), with particular organizational flexibility in lateral temporal regions during auditory processing and speech perception (Andric, Goldin-Meadow, Small, & Hasson, 2016; Andric & Hasson, 2015). That is, peripheral networks are not static but are, rather, dynamically organizing, e.g., as a function of context (Jeremy I. Skipper, 2015).

### 4.5. Conclusions

We have provided evidence for a core speech circuit comprised of primary motor, primary somatosensory, and primary auditory cortices that might be somewhat unique to humans based on connectivity data. This circuit manifests in specific connectivity patterns between the CS and TTG that are not evident for nearby auditory association regions in the STP that may explain why it has not often been documented in prior work: It relies on a relatively fine scale parcellation not used in the vast majority of studies. This CS-TTG circuit is speech associated and expresses genes highly conserved in vocal learning animals and related to prediction and efference copy. Thus, given the early ontological development of this circuit, we suggest that it serves as a basis for a vocal learning mechanism, namely, efference copy, that allows for maintenance and adjustment during production. As it is also active during perception, this same circuit could be reused to constrain interpretation of variable acoustic patterns arriving in auditory cortex (Jeremy I. Skipper et al., 2017; J. I. Skipper et al., 2006). More generally, this speech core is not well accounted for by current neurobiological models. We suggest that it could serve as an anchoring foundation for surrounding cortex in a core-periphery complex network organization. Integration of this perspective into a dynamic conception of the neurobiology of speech perception and language comprehension is an important goal for future work.

## Acknowledgments

Resting state data were provided by the Human Connectome Project, WU-Minn Consortium (Principal Investigators: David Van Essen and Kamil Ugurbil; 1U54MH091657). JIS would like to acknowledge support from Lily, NIH-NICHD K99/R00 HD060307 - “Neurobiology of Speech Perception in Real-World Contexts”, and EPSRC EP/M026965/1 “New pathways to hearing: A multisensory noise reducing and palate based sensory substitution device for speech perception.” UH’s work was conducted in part while serving at and with support of the National Science Foundation. Any opinions, findings, and conclusions or recommendations expressed in this material are those of the author(s) and do not necessarily reflect the views of the NSF.

